# Structural basis of cyclic phytocytokine recognition by the HSL3/NUT receptor

**DOI:** 10.64898/2025.12.22.696113

**Authors:** Hoseong Jung, Sejin Choi, Hyeonmin Ryu, Youngran Kim, Nari Seo, Yong Heum Na, Hyun Joo An, Kerstin Dürr, Kunhee An, Jong Hum Kim, Du-Hwa Lee, Bon-Kyoung Koo, Ji-Hye Yun, Ho-Seok Lee

## Abstract

Plant receptor kinases perceive diverse peptide signals to coordinate stress responses and developmental programs. The HAESA-LIKE 3 (HSL3/NUT) receptor recognizes CTNIP/SCREW phytocytokines—disulfide-stabilized cyclic peptides that regulate immune signaling and stress adaptation. However, how HSL3 distinguishes these structurally constrained cyclic peptides from linear signaling molecules remains unknown. Here we report near-atomic resolution cryo-EM structures of HSL3 in apo and CTNIP4^48-70^-bound states at ∼2.6 Å, using Arabidopsis CTNIP4 as a representative family member, revealing distinct mechanisms for cyclic peptide recognition. The conserved CTNIP motif occupies a negatively charged pocket in HSL3’s C-terminal region through a combination of polar contacts, hydrogen bonds, salt bridges, and van der Waals interactions. The receptor employs a two-step recognition mechanism—electrostatic steering followed by motif anchoring—that enables rapid ligand capture and release, consistent with the transient nature of stress signaling. Notably, an N-glycan at Asn449 directly contacts the CTNIP4 peptide, establishing glycosylation as an active participant in ligand recognition. Structure-guided mutagenesis combined with reactive oxygen species (ROS) burst assays confirmed the functional importance of key binding interfaces. N-terminal truncation experiments revealed a minimal active fragment: CTNIP4^51–70^ supported both rapid ROS production and sustained seedling growth inhibition, whereas the shorter CTNIP4^54–70^ variant retained ROS activity but failed to trigger long-term seedling growth inhibition. Structure-guided coevolutionary analysis across plant lineages reveals patterns of both conserved and variable receptor–ligand interfaces, highlighting evolutionary flexibility while preserving core features of recognition. These conserved recognition principles, mediated by receptor glycosylation and evolutionary plasticity, enable specificity in peptide signaling with implications for engineering stress-resilient crops.

## Introduction

Plants continuously monitor their extracellular environment to adjust growth and activate stress responses, relying on cell-surface receptor kinases that translate outside signals into intracellular decisions (Jian et al., 2024; Lee et al., 2021). Among these, leucine-rich repeat receptor kinases (LRR-RKs) form one of the largest signaling modules in plants and are deeply embedded in coordinating development, immunity, and environmental adaptation (Dievart et al., 2020; Ryu et al., 2025). Their ability to discriminate among diverse endogenous and exogenous cues is central to how plants maintain resilience under fluctuating conditions (Choi and Lee, 2024; Furumizu and Aalen, 2023).

Structural studies of plant peptide-receptor complexes, including the IDA-HAESA pair, have focused primarily on linear peptide ligands that engage receptors through extended conformations (Roman et al., 2022; Santiago et al., 2016; Zhang et al., 2016). However, how LRR-RKs recognize peptides with constrained, disulfide-stabilized folds—an architecture increasingly associated with stress signaling—remains unexplored.

The HSL3/NUT receptor provides a compelling case. This receptor recognizes CTNIP/SCREW peptides, a family of stress-responsive ligands whose disulfide-constrained structure contrasts with these previously characterized linear ligands. CTNIP expression is rapidly elevated by drought and biotic stresses, and their perception by HSL3 initiates a BAK1-dependent signaling pathway that triggers ROS production and seedling growth inhibition (Bjornson et al., 2021; Li et al., 2014; Liu et al., 2022; Rhodes et al., 2022b). How HSL3 accommodates this constrained peptide architecture—and achieves specificity for CTNIPs over other endogenous signals—has remained unknown. In this study, we use Arabidopsis CTNIP4 as a representative CTNIP family member and a CTNIP4-derived core fragment as the ligand to define the structural basis of HSL3 recognition.

Adding further complexity, LRR-RK ectodomains are extensively N-glycosylated (Haweker et al., 2010; Nagashima et al., 2018). These glycans are generally assumed to assist folding, maturation, and ER quality control, yet emerging evidence suggests that specific glycan moieties can contribute directly to ligand engagement and co-receptor recruitment (Jia et al., 2024; Wu et al., 2024). Whether such glycan-mediated tuning plays a role in HSL3 signaling, particularly given the unusual disulfide-stabilized architecture of CTNIPs, has not been resolved.

Here, we present near-atomic-resolution cryo-electron microscopy structures of HSL3 in both apo and CTNIP4^48–70^-bound states, integrated with biochemical binding measurements, structure-guided mutagenesis, AlphaFold modeling, and in-plant functional analyses. Our results reveal how the conserved CTNIP C–T–N–I–P motif is precisely anchored within a C-terminal pocket of HSL3 and demonstrate that a strategically positioned N-glycan at Asn449 dynamically stabilizes peptide accommodation. Structure-guided coevolutionary analysis across plant lineages further reveals that receptor–ligand interfaces exhibit both highly conserved interaction hotspots and selectively variable surfaces, suggesting that this signaling module balances core recognition with lineage-specific tuning. This unexpected glycan contribution, coupled with evolutionary plasticity at the binding interface, illustrates previously unrecognized layers of specificity within LRR-RK signaling and establishes a structural mechanism by which receptors can engage disulfide-stabilized cyclic-like peptides.

Together, our findings define the molecular logic of the HSL3–CTNIP4 module and highlight how receptor glycosylation, pocket architecture, and evolutionary diversification together expand the signaling repertoire of plant receptor kinases. This work provides a foundational framework for understanding how plants diversify peptide recognition to fine-tune immune and stress responses.

## Results

### Overall architecture of HSL3 revealed by high-resolution cryo-EM

The ectodomain of *Arabidopsis thaliana* HAESA-LIKE 3 (HSL3) was recombinantly expressed, purified to homogeneity, and subjected to single-particle cryo-electron microscopy. The receptor eluted as a single symmetric peak during size-exclusion chromatography and appeared as a sharp band in SDS–PAGE and anti-His immunoblotting, confirming its high purity and monodispersity (Supplemental Figure 1). Cryo-EM data collection and 3D reconstruction yielded near-atomic-resolution maps for both apo HSL3 (2.63 Å) and the CTNIP4^48–70^-bound complex (2.59 Å) (Supplemental Figures 2 and 3). Model–map agreement was consistently high (Mask CC = 0.74–0.66), and both structures exhibited excellent stereochemical quality, with MolProbity scores of 2.53 and 2.82, respectively (Supplementary Table 1).

**Figure 1.**
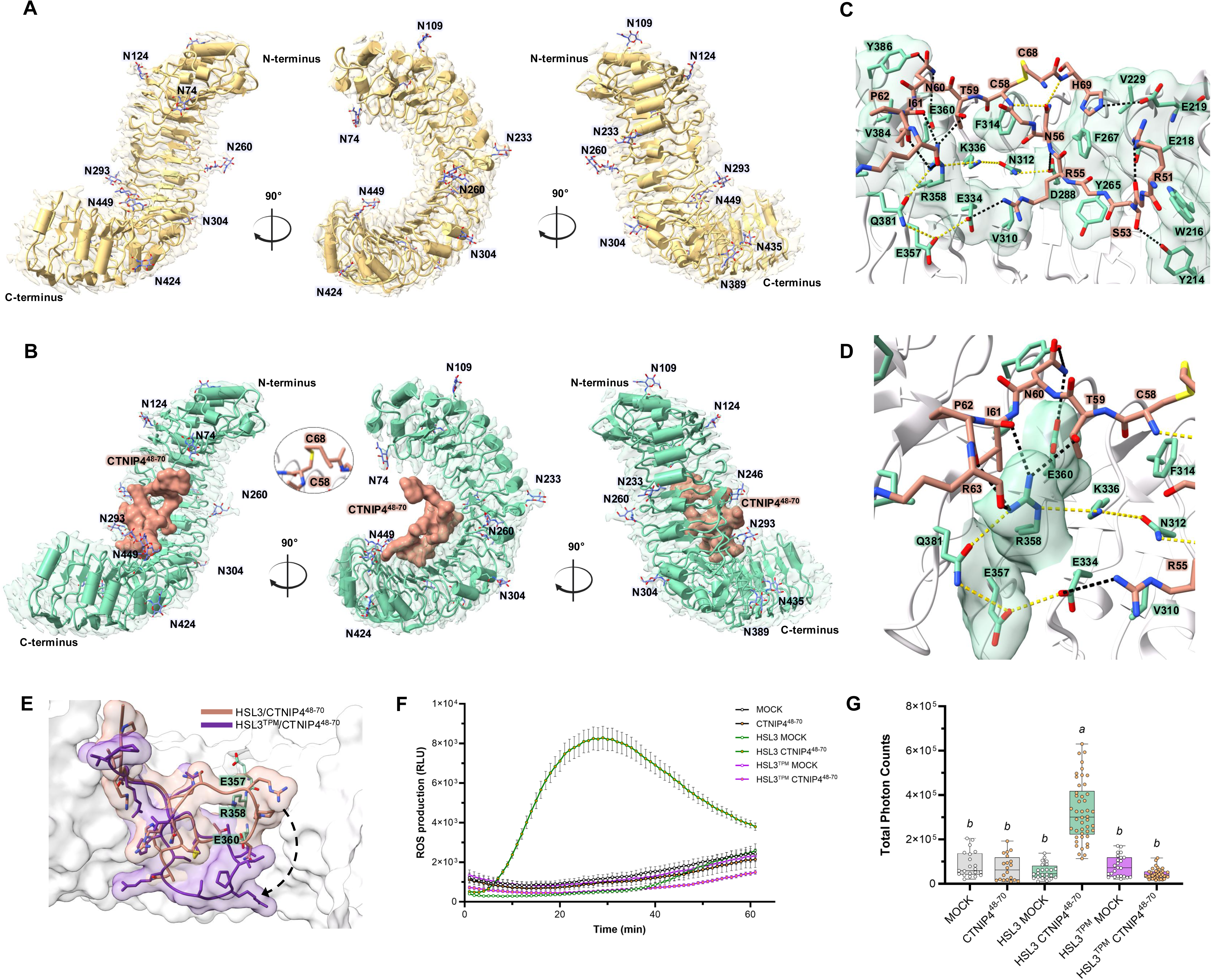
Cryo-EM structures of HSL3–CTNIP4 complexes reveal a unique recognition mode mediated by disulfide-constrained cyclic peptide. **(A)** Cryo-EM structure of apo HSL3 is displayed at 2.63 Å resolution: The ectodomain adopts a right-handed solenoid LRR architecture with well-defined N- and C-terminal caps. The cryo-EM density is shown as a semi-transparent surface, overlaid with the refined atomic model rendered in light golden-beige ribbon representation. N-linked glycans, visualized as stick models and colored in vivid periwinkle blue, are labeled at their respective glycosylation sites (Asn74, Asn109, Asn124, Asn233, Asn246, Asn260, Asn293, Asn304, Asn389, Asn424, Asn435, and Asn449). Three orthogonal views (90° rotations) highlight the distribution of glycans along the inner and outer faces of the LRR solenoid. **(B)** Cryo-EM structure of the HSL3–CTNIP4 complex at 2.59 Å resolution. The HSL3 ectodomain (mint green; ribbon) engages the CTNIP4 peptide (residues 48–70; soft terracotta; surface) within a concave pocket located at the C-terminal end of the LRR solenoid. The peptide adopts a disulfide-constrained cyclic conformation, stabilized by the Cys58–Cys68 linkage (highlighted in the inset), which supports the compact loop architecture required for binding. N-linked glycans and their attachment sites are labeled as in panel (A). **(C)** Detailed view of the HSL3–CTNIP4 interface. The CTNIP4 peptide (soft terracotta) and HSL3 (mint green) are shown in stick representation for all residues located within 4 Å of the interface. Direct receptor–peptide interactions—including hydrogen bonds, salt bridges, and polar contacts—are depicted as black dashed lines, highlighting the primary stabilizing network that anchors the ligand within the binding pocket. Internal hydrogen bonds and polar interactions within CTNIP4 or within HSL3 that indirectly support peptide accommodation are shown as yellow dashed lines, emphasizing their role in stabilizing the local structural environment around the interface. Residues involved in non-bonded van der Waals contacts are also shown as sticks and labeled accordingly, providing a complete map of intermolecular contacts shaping the recognition surface. **(D)** Close-up view of the electrostatic cluster within the HSL3–CTNIP4 interface. The CTNIP4 peptide (soft terracotta) and HSL3 (mint green) are shown with interacting residues within 4 Å represented as sticks. A linear electrostatic cluster formed by residues Glu357, Arg358, and Glu360 lines the inner surface of the binding groove. Direct peptide–receptor interactions—including hydrogen bonds, salt bridges, and polar contacts—are depicted as black dashed lines. Yellow dashed lines indicate intramolecular hydrogen bonds or polar contacts within HSL3 or CTNIP4 that contribute indirectly to stabilizing the binding environment by reinforcing the local structural framework. Together, these interactions outline the electrostatic hub centered on E357–R358–E360 that shapes the ligand-binding pocket. **(E)** Comparison of CTNIP4 binding modes between *Arabidopsis thaliana* wild-type HSL3 and the triple-mutant receptor. Binding conformations and positions of CTNIP4 are compared between wild-type HSL3 (CTNIP4 shown in soft terracotta) and an AlphaFold3-predicted triple mutant (E357A/R358A/E360A, HSL3^TPM^; CTNIP4 shown in royal purple). In both structures, the N-terminal portion of the peptide (residues 48–59) aligns well; however, in the mutant complex, the C-terminal region (residues Asn60–Asn70) bends outward, resulting in a marked shift in binding position. The three mutated residues within HSL3 are highlighted as mint-green stick models. Dashed outlines indicate the direction of peptide displacement relative to the wild-type binding mode. **(F)** Reactive oxygen species (ROS) burst profiles in *Nicotiana benthamiana* leaves transiently expressing AtHSL3 or AtHSL3^TPM^. Leaf discs were treated with 1 µM CTNIP4^48-70^ or MOCK (water) and chemiluminescence was recorded for 60 min. Data represent means ± SEM from biologically independent samples for each condition. **(G)** Total photon counts (area under the curve) were measured over 60 min from the same dataset shown in Figure 1F. Box plots represent the first and third quartiles centered on the median, with whiskers showing the minimum-maximum range. Individual datapoints indicate biologically independent observations. Statistical differences among groups were assessed using one-way ANOVA with Tukey’s multiple comparison test (*P* < 0.05). Distinct letters denote statistically different groups at the 95% confidence level.

**Figure 2.**
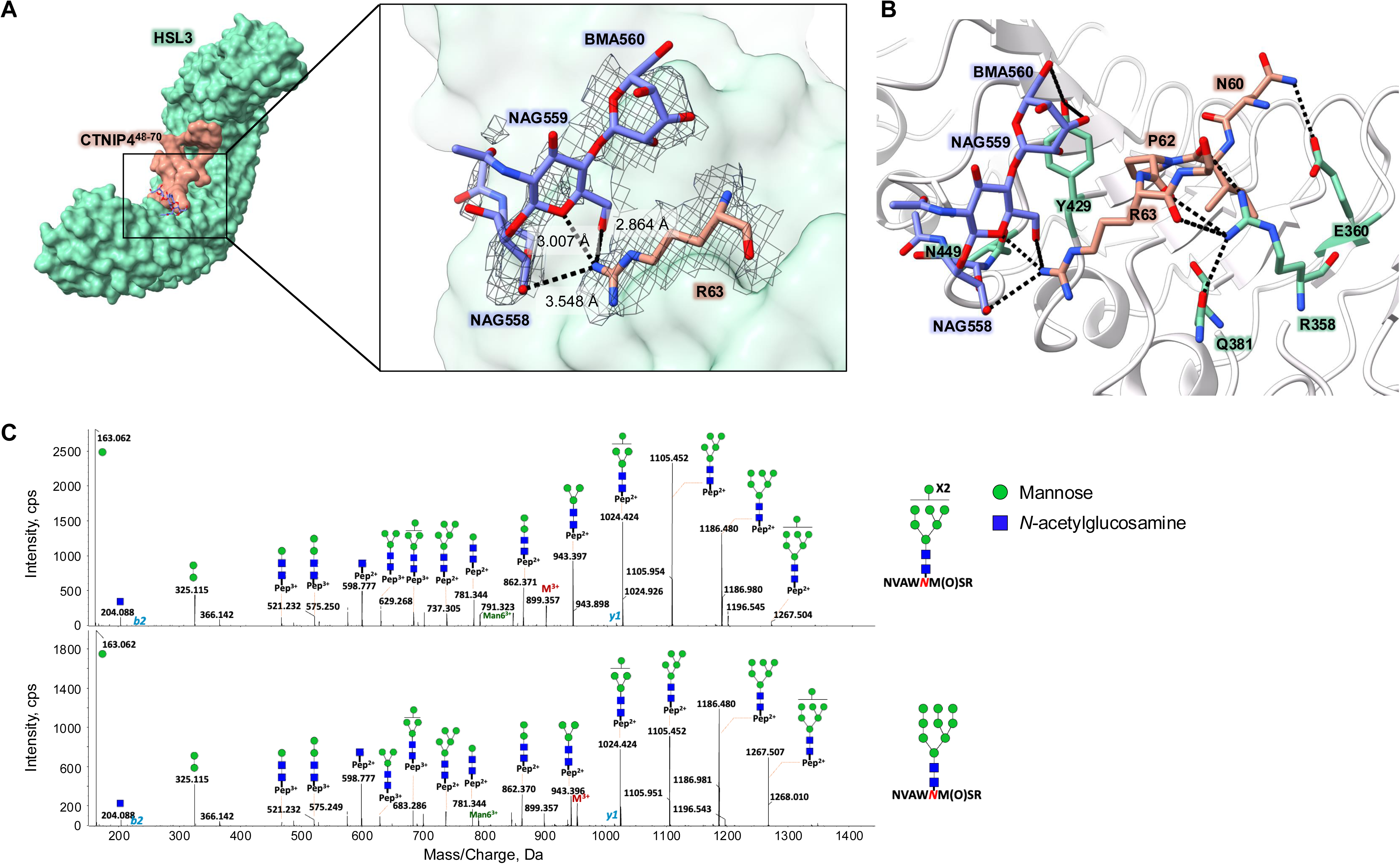
Glycan-mediated interactions within the HSL3–CTNIP4 binding interface. (**A**) Detailed views of the HSL3-CTNIP4 interface near the conserved CTNIP motif along with a glycan. The Asn449-linked N-glycan (vivid periwinkle blue; stick models) and the interacting CTNIP4 residue Arg63 (soft terracotta; stick models) are shown together with the HSL3 surface (mint green). The cryo-EM local density is displayed as a grey mesh contoured at an appropriate threshold. Dashed black lines indicate measured hydrogen bonds or polar contacts, and corresponding distances are labeled. (**B**) Enlarged view of the glycan–peptide contact region. NAG559 forms a hydrogen bond with CTNIP4 Arg63, while BMA560 interacts with HSL3 Tyr429. Additional receptor–peptide contacts—such as interactions involving Arg358 and Glu360—are indicated with black dashed lines. Glycan residues are shown in vivid periwinkle blue, CTNIP4 in soft terracotta, and HSL3 side chains in mint green stick representation. **(C)** Identification of N-glycosylation at Asn449 on HSL3. Collision-induced dissociation (CID) MS/MS spectra of the N-glycopeptide NVAWNM(O)SR carrying Man_8_GlcNAc₂ at *m*/*z* 899.357 ([M + 3H]^3+^) (upper) and Man_9_GlcNAc₂ at *m*/*z* 953.345 ([M + 3H]^3+^) (lower) from HSL3. For the glycan cartoons, green circles denote mannose and blue squares denote *N*-acetylglucosamine.

**Figure 3.**
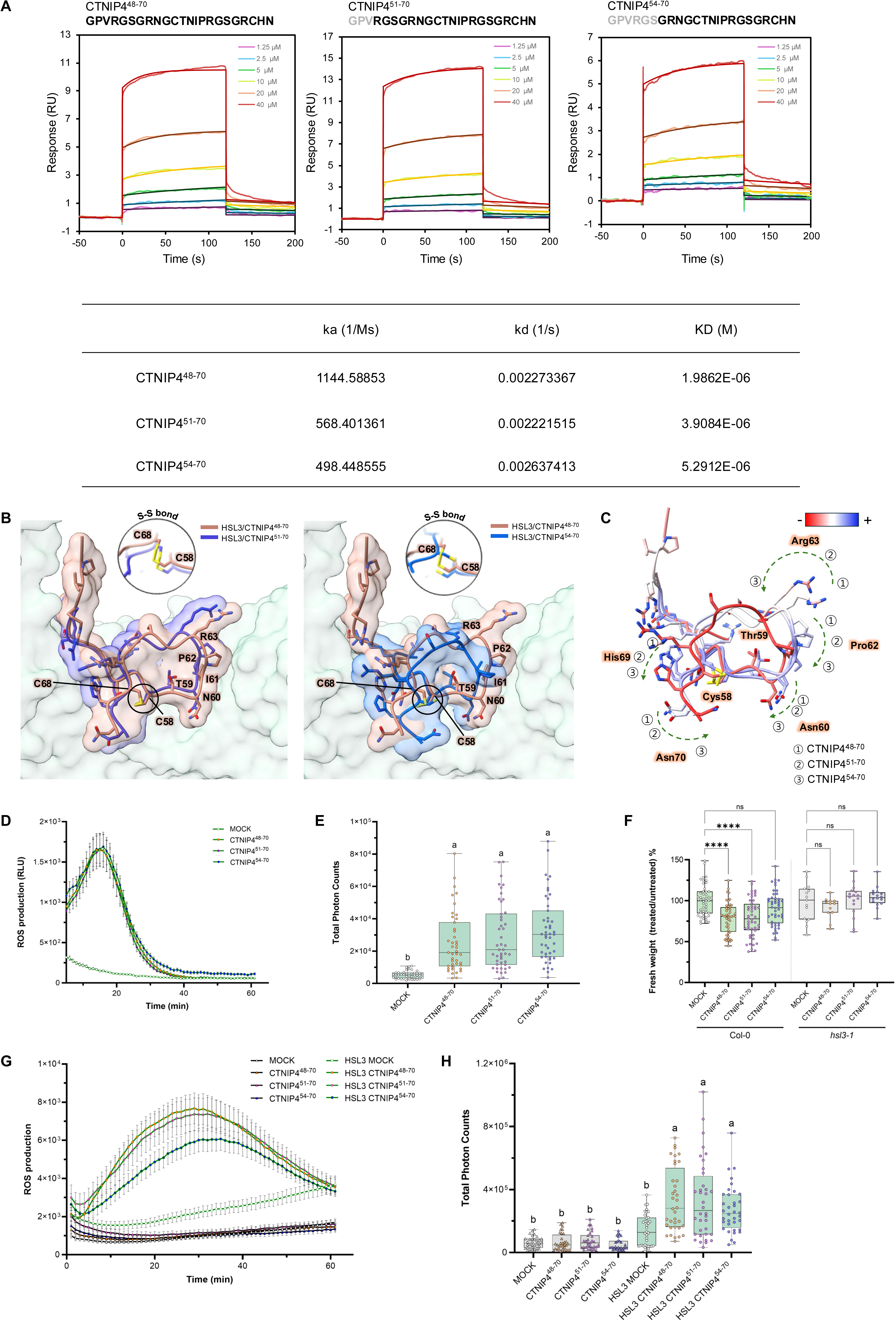
Truncation and modeling define the minimal CTNIP4 length required for HSL3 binding and signaling. **(A)** Surface plasmon resonance (SPR) sensorgrams for CTNIP4^48–70^, CTNIP4^51–70^, and CTNIP4^54–70^. Each panel shows a concentration series (1.25, 2.5, 5, 10, 20, 40 μM) with experimental sensorgrams (solid lines) and global 1:1 Langmuir fitting curves shown in the same colors. Vertical lines indicate the association and dissociation phases corresponding to the start of analyte injection and the transition to buffer flow, respectively. Kinetic parameters (k_a_, k_d_, KD) are listed below each panel. **(B)** Comparison of HSL3-bound CTNIP4 peptide models of different lengths. Superimposed structural models of CTNIP4 variants bound to HSL3 are shown. Left panel: CTNIP4^48–70^ (soft terracotta) compared with CTNIP4^51–70^ (deep indigo blue). Right panel: CTNIP4^48–70^ (soft terracotta) compared with CTNIP4^54–70^ (vibrant cobalt blue). The HSL3 receptor surface is displayed in mint green, and core residues of CTNIP4 are labeled for reference. The Cys58–Cys68 disulfide bond that stabilizes the cyclic conformation of CTNIP4, as well as its presence or absence across peptide-length variants, is highlighted in the inset to illustrate how ligand truncation affects S–S bond formation. **(C)** B-factor–mapped structural comparison of CTNIP4 variants bound to HSL3. Superimposed models of CTNIP4^48–70^, CTNIP4^51–70^, and CTNIP4^54–70^illustrate structural variation among the three peptides upon N-terminal truncation. For each peptide, B-factors are color-coded from low (blue) to high (red), highlighting regions of differential flexibility. The Cys58–Cys68 disulfide bond is depicted as yellow sticks. **(D)** ROS burst profiles in *Arabidopsis thaliana* leaf discs treated with 1 µM N-terminally truncated CTNIP4 variants (CTNIP4^48-70^, CTNIP4^51-70^, CTNIP4^54-70^) or MOCK (water). The initial 0-5 min interval was excluded as background noise. Data represent means ± SEM from biologically independent samples. **(E)** Total photon counts (area under the curve over 5-60 min) calculated from the dataset shown in Figure 3D. Statistical analysis and box-plot formatting were performed as described for Figure 1G. Distinct letters denote significantly different groups. **(F)** Fresh weight of *Arabidopsis* seedlings treated with 5 µM N-terminally truncated CTNIP4 variants or MOCK (water) for 7 days, expressed as percentage relative to the MOCK control within each genotype. Statistical analysis and box-plot formatting were performed as described for Figure 1G. Each point represents an individual seedling. Asterisks indicate significance relative to the corresponding MOCK control (*P* < 0.0001; ns, not significant). **(G)** ROS responses to N-terminally truncated CTNIP4 variants in HSL3-expressing *N. benthamiana* leaves. Leaf discs were treated with each elicitor or MOCK (water) at a final concentration of 1 µM. Data represent means ± SEM from biologically independent samples. **(H)** Total photon counts (area under the curve over 0-60 min) calculated from the same dataset shown in Figure 3G. Statistical analysis and box-plot formatting were performed as described for Figure 1G. Distinct letters denote significantly different groups.

HSL3 adopts a canonical horseshoe-shaped LRR architecture composed of 21 leucine-rich repeats flanked by well-defined N- and C-terminal capping domains (Figure 1A and 1B). Its overall scaffold is reminiscent of the floral abscission receptor HAESA (Santiago *et al*., 2016) and structurally related LRR-RKs, yet HSL3 displays distinct surface electrostatics across the C-terminal repeats that give rise to a uniquely contoured ligand-binding environment. The central solenoid forms a continuous parallel β-sheet supported by inward-facing hydrophobic cores, whereas the convex outer surface is decorated with multiple glycan densities, consistent with previous observations from other LRR-RK ectodomains (Jia *et al*., 2024; Wu *et al*., 2024).

The N-terminal “hook” region contains two disulfide-bonded cysteine pairs (Cys53–Cys60 and Cys84–Cys110) that secure the proximal repeats into a compact arrangement, effectively reinforcing the solenoid core. In contrast to HAESA—which harbors additional cysteines in its C-terminal cap—HSL3 contains a single disulfide linkage between Cys375 and Cys401 positioned adjacent to the peptide-binding surface, where it locally enhances rigidity and shapes the ligand-recognition channel. The well-resolved loop and side-chain densities enabled reliable modeling of the disulfide-stabilized motifs and allowed clear visualization of the conserved asparagine ladder along the inner β-sheet, which supports the stability of the LRR scaffold.

### CTNIP4 binding rigidifies the local pocket and induces subtle structural rearrangements

The CTNIP448–70 peptide is unambiguously resolved in the cryo-EM density, occupying a concave pocket formed by the C-terminal LRRs of HSL3 (Supplemental Figure 4). The peptide adopts a compact, ring-like conformation stabilized by an internal Cys58–Cys68 disulfide bond—an architectural feature that distinguishes CTNIPs from previously characterized linear LRR-RK ligands such as IDA or CLE peptides (Hohmann et al., 2017; Santiago *et al*., 2016). The conserved CTNIP4 core motif (C–T–N–I–P) is precisely anchored within the pocket through an extended network of hydrogen bonds and polar interactions formed by HSL3 residues Arg 358, Glu360 and Tyr386. Additional stabilization is provided by positively charged CTNIP4 residues Arg51, Arg55, and His69, which complement the acidic binding groove of HSL3, while Ser53 and Asn56 contribute additional polar contacts that further reinforce the interface (Figure 1C and Supplemental Table 2).

The N-terminal segment of the peptide (Gly48–Asn56), together with the C-terminal His69, lies across a negatively charged surface formed by HSL3 residues Glu218, Glu219, Asp288, and Glu334, establishing multiple salt bridges that orient the ligand for precise pocket engagement. Arg51, Arg55, and His69 form salt bridges with Glu218, Glu219, and Glu334, respectively, while Ser53 hydrogen-bonds with Tyr214 and Asn56 engages Asp288 through a polar contact that stabilizes the N-terminal arm. Within the conserved motif, Asn60, Ile61, Pro62, and Arg63 form additional polar and hydrogen-bonding interactions with Glu360, Arg358, and Tyr386, further reinforcing the orientation of the cyclic loop (Figure 1C and Supplemental Table 2).

Electrostatic surface potential calculations revealed that the concave face of HSL3 forms a broadly acidic landscape, which complements the positively charged patches along the CTNIP4 peptide (Supplemental Figure 5A). Upon binding, the N-terminal residues Gly48–Arg55 rest over an acidic depression, while the disulfide-stabilized CTNIP motif inserts into a hydrophobic groove at the core of the binding site (Supplemental Figure 5B). This arrangement establishes a two-layered recognition mechanism combining electrostatic steering at the entrance of the pocket with hydrophobic readout at its core, ensuring rapid yet selective capture of the cyclic ligand.

Superposition of apo and CTNIP4-bound HSL3, followed by residue-wise r-DDM analysis (Apo – Complex), revealed a nonuniform pattern of structural adjustments across the ectodomain (Supplemental Figure 6). Residues in the N-terminal portion of the binding interface (approximately residues 210–280) exhibited predominantly negative values, indicating that these regions shift outward upon ligand engagement. In contrast, residues forming the core CTNIP4-binding groove (roughly residues 280–390) showed mainly positive differences, consistent with a local compaction that accompanies insertion of the cyclic CTNIP motif. This spatially graded pattern suggests that CTNIP4 binding expands the entrance of the pocket while tightening the central groove to stably accommodate the disulfide-stabilized peptide. Regions distal to the binding site—including portions of the N-terminal LRRs and the terminal repeats (approximately residues 400–540)—displayed a mixture of positive and negative differences, reflecting distributed long-range adjustments rather than a strictly localized conformational change. Consistent with this behavior, ΔB-factor analysis (Complex – Apo) showed an overall reduction in B-factors across the ectodomain, with the most pronounced decreases in distal regions, whereas the central binding site exhibited more modest stabilization. Together, these analyses indicate that CTNIP4 engagement induces global rigidification of HSL3 while enabling local structural fine-tuning necessary for accommodating the cyclic ligand.

### Mutational and functional characterization identify structure−function relationship between the electrostatic cluster and peptide orientation

As mentioned above, Arg358 and Glu360 are essential residues binding CTNIP4 peptide through direct hydrogen bonds, thus making CTNIP4 anchoring and orientation determinants. Although Glu357 itself has does not directly contact the peptide, it lies beneath Arg358 and Glu360, where it interacts with neighboring residues to maintain the structure of the binding pocket in space. These three residues are positioned in a linear fashion along the inner surface of the binding groove, and together they constitute an electrostatic cluster that molds the local potential landscape of critical importance for CTNIP4 recognition (Figure 1D). When these three residues were mutated to alanine (HSL3TPM, E357A/R358A/E360A), AlphaFold3 predicted a marked distortion in the CTNIP4 backbone from Asn60. In this mutant, an increase is observed in the distance between Arg63 and Asn449-linked N-glycan, as well as in the distortion of the peptide loop relative to the wild-type complex (Figure 1E). In agreement with these structural predictions, ROS burst assays performed in Arabidopsis lines expressing the HSL3TPM forms showed significantly decreased CTNIP4-induced ROS bursts (Figure 1F and 1G), whereas total protein expressions were not affected (Supplemental Figure 7). Altogether, these results not only support the structural model of both forms but also underscore the functional importance of the Glu357–Arg358–Glu360 electrostatic cluster in keeping the peptide properly oriented as well as mediating a stable HSL3−CTNIP4 association. This cluster therefore serves as an essential connector between ligand binding and receptor activation, allowing fidelity of signal transduction.

### Glycan-assisted stabilization of the peptide–receptor interface

Analysis of the HSL3 ectodomain structure by cryo-EM demonstrated that glycosylation is present, on twelve asparagine residues from within the LRR scaffold (Figure 1A and 1B). To determine the overall composition of these glycans, we performed global N-glycan profiling using MALDI–TOF mass spectrometry, which identified two dominant glycoforms—HexNAc(8)Hex(2) and HexNAc(9)Hex(2)—along with seven minor variants (Supplemental Figure 8). In the cryo-EM map, strong local density is observed for the Asn449-linked N-glycan, resolved as a branched NAG–NAG–BMA tri-saccharide (NAG558, NAG559, and BMA560). The middle GlcNAc unit (NAG559) extends toward the bound CTNIP4 peptide and forms a direct interaction with Arg63, establishing a glycan-assisted contact at the binding interface (Figure 2A and 2B). The guanidinium group of Arg63 is positioned within hydrogen-bonding distance of the NAG559 O6 and O7 atoms (2.86 Å and 3.00 Å, respectively), suggesting the presence of hydrogen bonds or electrostatic contacts. Although Arg63 exhibits a relatively elevated B-factor (∼36 Å², compared with the average of ∼22 Å²), the interaction appears to contribute to local stabilization of the interface without fully restricting the intrinsic conformational mobility of the cyclic CTNIP4 peptide. Additionally, BMA560 forms a direct interaction with HSL3 Tyr429, completing a glycan-mediated bridging network between the receptor and the peptide. This interaction further reinforces the local stabilization at the HSL3–CTNIP4 interface. MALDI-TOF mass spectroscopy of the glycan structure at Asn449 indicated that two major glycoforms, HexNAc(8)Hex(2) and HexNAc(9)Hex(2), were present. This observation emphasizes the diversity of glycans at this site (Figure 2C). These interactions between receptor glycans and peptides are important for the structural integrity of the binding pocket and contribute to increased fidelity of ligand recognition. It is possible that this dynamic contact would cooperate to provide rapid on and off kinetic rates, which contribute to the structural flexibility for the reversible signaling (Haweker *et al*., 2010; Wu *et al*., 2024). This glycan-assisted stabilization is consistent with that observed in the MIK2–SCOOP–BAK1 complex (Jia *et al*., 2024; Wu *et al*., 2024). Here, it is just one example where glycosylation of receptors is not only critical for structural integrity but also involved in ligand recognition and coreceptor binding. By contrast, in HSL3, the Asn449-bound glycan is shown to be involved directly into interaction with the CTNIP4 peptide. This demonstrates the active role of receptor glycosylation in modulating ligand binding and increasing receptor stability.

### N-terminal truncation establishes the minimal CTNIP motif necessary for HSL3 recognition and signaling

To map the minimal binding region required for HSL3 association, we performed Surface Plasmon Resonance (SPR) using three N-terminally truncated variants of CTNIP: CTNIP4^48–70^, CTNIP4^51–70^ and CTNIP4^54–70^ (Figure 3A). All peptides bound to HSL3, but with progressively reduced affinities (KD =1.99 × 10⁻⁶, 3.91 ×10⁻⁶ and 5.29 × 10⁻⁶ M, respectively). The dissociation rate constant (K_d_) showed minimal variation among the variants, whereas the association rate constant (K_a_) decreased about twofold, from 1.14 × 10³ to 5.68 × 10² and 4.98 × 10² M⁻¹·s⁻¹, respectively. This kinetic profile indicates that truncation of the N-terminal residues does not a destabilizing factor the final complex; instead, it primarily impairs the initial association rate between peptide and receptor. Mechanistically, the N-terminal residues (Gly48–Arg55) are predicted to be critically involved in electrostatic steering and hydrogen bonding, which guide the peptide into an optimal orientation for binding to HSL3’s acidic groove. Once the conserved CTNIP motif (C–T–N–I–P) is correctly engaged, the dissociation kinetics remain similar, explaining the nearly constant K_d_ values. Accordingly, the increased KD values largely reflect a decrease in K_a_ with little or no change in complex stability.

Although the truncated CTNIP4 peptides used for SPR retain the Cys58–Cys68 disulfide bond, their binding affinities decreased markedly upon N-terminal shortening. Consistent with this, AlphaFold3 modeling of the corresponding truncated sequences shows that removal of N-terminal residues destabilizes the overall cyclic fold, often leading to disruption of the Cys58–Cys68 disulfide constraint. Models in which the S–S bond is not preserved fail to insert the core motif properly into the HSL3 pocket, suggesting that the N-terminal region contributes not only to engagement of the acidic groove but also to maintaining the disulfide-stabilized geometry required for precise binding (Figure 3B).

The CTNIP motif region exhibited consistently low B-factor values, indicating a structurally rigid core of interaction, whereas the peripheral residues displayed increased flexibility. The relative positioning of the key residues Cys58 and Thr59 remained largely invariant across all variants, while residues above the motif—Asn60, Pro62, His69, and Asn70—underwent gradual inward displacement toward the receptor surface with increasing truncation (Figure 3C). In the CTNIP4^54–70^, disruption of the Cys58–Cys68 disulfide bond caused a pronounced relocation of Arg63 and elevated B-factors for His69 and Asn70, reflecting local destabilization and loss of structural restraint. These conformational changes demonstrate that the disulfide linkage and N-terminal anchoring cooperatively maintain the correct topology of the CTNIP4 loop necessary for stable HSL3 recognition.

To functionally validate these structural insights, we performed in planta assays which revealed that N-terminal truncation progressively diminishes CTNIP4 function, with differential effects on distinct physiological responses. When applied at 1 µM, all three CTNIP4 variants elicited robust and comparable ROS bursts in *Arabidopsis*, with total photon counts separated from the MOCK controls (Figure 3D and 3E). These data indicate that early HSL3-dependent signaling events tolerate substantial N-terminal shortening as long as the conserved C–T–N–I–P core remains intact, consistent with the preserved dissociation kinetics observed in the SPR assays. However, long-term physiological outputs revealed a stricter requirement for the N-terminal segment. In the seedling growth-inhibition assay, performed using 5 µM of the peptide, CTNIP4^48–70^ and CTNIP4^51–70^ markedly reduced fresh weight, whereas CTNIP4^54–70^ failed to induce any measurable inhibition and was indistinguishable from the mock treatment (Figure 3F). Notably, all three variants lost growth-inhibitory activity in the *hsl3-1* background, confirming that these responses are strictly HSL3-dependent. This divergence between early ROS production and long-term growth inhibition suggests that low-affinity or less-stable ligand conformations can still trigger transient early signaling but are insufficient to support the sustained or cumulative signaling required for growth repression. A comparable trend was noted in *N. benthamiana*, where CTNIP4 variants applied at 1 µM failed to trigger ROS production unless HSL3 was heterologously expressed, confirming strict receptor dependence. In this system, CTNIP4^54–70^ elicited a noticeably weaker ROS peak compared with the longer variants, although total photon counts still clustered closely (Figure 3G and 3H). These results suggest that progressive N-terminal truncation subtly reduces signaling potency, consistent with reduced binding efficiency observed in vitro.

Together, these results demonstrate that while the minimal CTNIP motif is sufficient for initiating rapid immune outputs, productive long-term signaling requires the N-terminal anchoring residues that stabilize the disulfide-constrained fold and enhance receptor engagement. Accordingly, the minimal CTNIP4 unit capable of sustaining both ligand activity and receptor binding affinity must extend beyond residues 54–70, emphasizing the importance of the N-terminal segment in ensuring productive interaction.

### Core Conservation and Peripheral Remodeling Define the Evolution of the HSL3–CTNIP4 Signaling Interface

To elucidate how the HSL3–CTNIP4 recognition system has evolved across land plants, we integrated residue-level conservation maps, expanded interface analyses, residue-pair entropy measurements, and structure-based similarity clustering (Figure 4 and Supplemental Figures 9–12). Collectively, these data reveal that the HSL3–CTNIP4 interface follows a dual evolutionary pattern in which a deeply conserved receptor core supports lineage-specific diversification of the surrounding interaction surface and its cognate peptide ligand.

**Figure 4.**
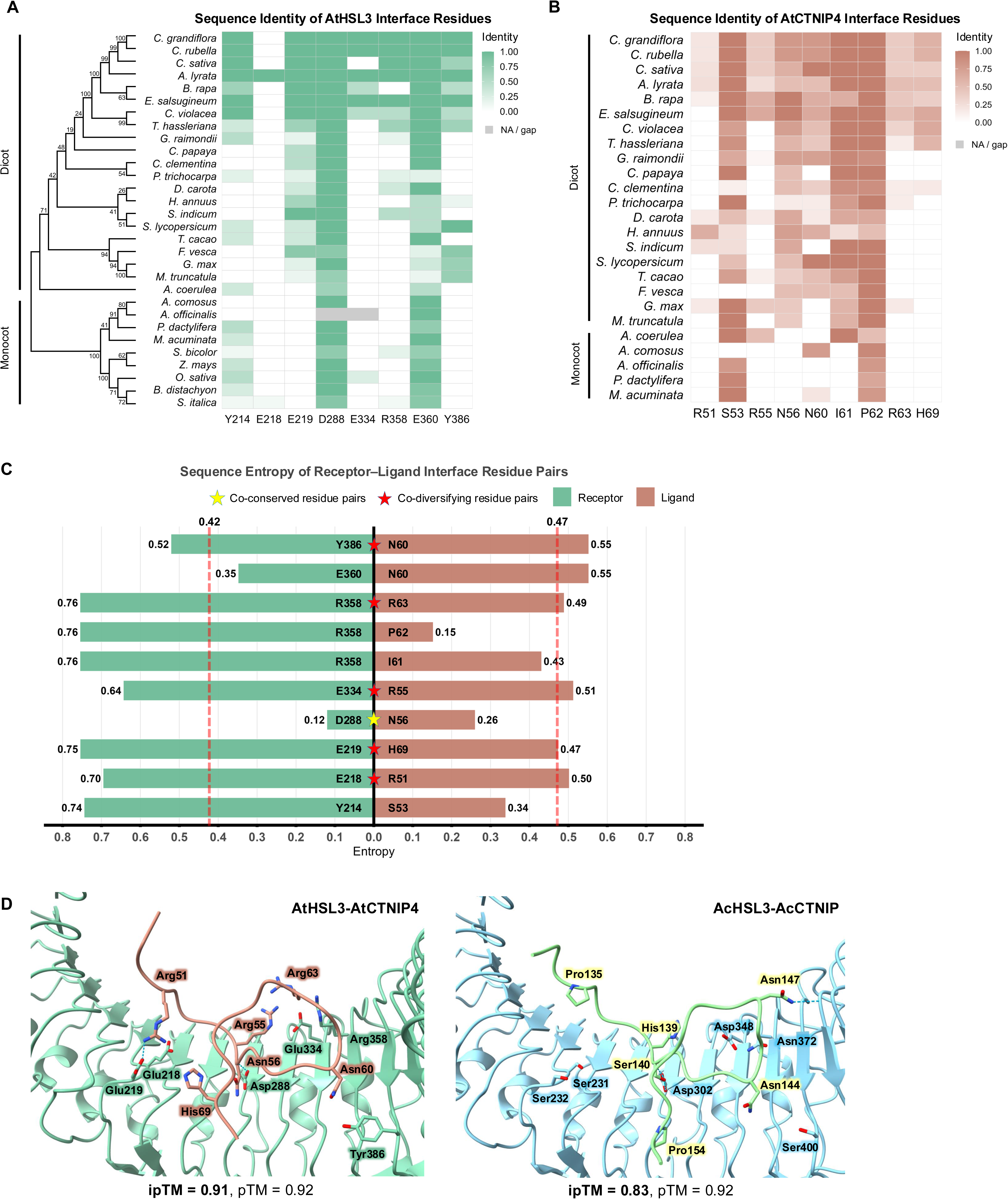
Structure-guided conservation and divergence at the HSL3–CTNIPs interface. **(A)** Conservation of bonding residues at the HSL3–CTNIPs interface (receptor side). Heatmap shows reference-based identity to AtHSL3 at eight ectodomain sites that make bonding interactions with CTNIP4 (hydrogen bond, salt bridge, or polar contact ≤ 4.0 Å): Y214, E218, E219, D288, E334, R358, E360, Y386 (AtHSL3 numbering). Identity was scored per homolog (1 if identical to AtHSL3, 0 otherwise; gaps = NA) and averaged per species (0–1). Rows are ordered by a maximum-likelihood guide tree (MEGA12) from the HSL3-ECD MSA; clades (monocots, dicots) are bracketed. Color scale: 0–1; gray, NA/gap. **(B)** Conservation of bonding residues at the HSL3–CTNIPs interface (ligand side). Heatmap of identity to AtCTNIP4 at nine CTNIP positions that form bonding contacts with HSL3 (AtCTNIP4 numbering): R51, S53, R55, N56, N60, I61, P62, R63, H69. Scoring and aggregation follow panel (A) (1/0 per homolog; species value = mean; gaps = NA). Species shown may differ from (A) because curated CTNIP homologs were unavailable or failed motif/length filters in some genomes. Color scale: 0–1; gray, NA/gap. **(C)** Sequence entropy of HSL3–CTNIPs bonding residue pairs. Bar plot of per-site sequence entropy for bonding interface pairs. Receptor entropies were computed from the HSL3 ectodomain MSA; ligand entropies from the CTNIP residues 37-70 alignment. For each set (receptor, ligand), the median entropy was used as the threshold (vertical dashed lines). Pairs with both entropy < median are classified as co-conserved (yellow star); pairs with both entropy ≥ median are co-diversifying (red star). Bars are labeled with entropy values; residue labels follow AtHSL3/AtCTNIP4 numbering. **(D)** Structural context of the bonding interface pairs. Left, AlphaFold3 model of the AtHSL3–AtCTNIP4 complex with the star-marked interface residues from (C) highlighted and labeled. Right, AlphaFold3 complex prediction for *Ananas comosus* HSL3 paired with a putative CTNIP-like candidate. Equivalent positions were selected by MSA mapping from the Arabidopsis interface (used for annotation/visualization only, not to redefine contacts).

Analysis of the eight bonding residues in HSL3 (Figure 4A) showed that while the core acidic anchor points (D288 and E360) are strictly conserved, the surrounding interface residues display lineage-specific divergence. Most dicots retain the canonical Arabidopsis-like signature, whereas both monocots and a subset of dicots exhibit substantial variation at these peripheral bonding positions. This trend was reinforced in the expanded interface analysis (Supplemental Figure 9A), which includes both bonding and non-bonded contacts: while the majority of residues vary across lineages, two acidic positions—D288 and E360—remain nearly invariant across all examined species, defining a stable electrostatic core essential for CTNIP4 binding. Sequence logos (Supplemental Figures 9B–D) further indicated that this receptor scaffold is evolutionarily constrained at core positions but flexible at outer surface residues, giving rise to distinct conserved and diversified subgroups within dicots.

In contrast to the receptor, the CTNIP ligand interface generally displays a higher degree of overall evolutionary remodeling, though key residues remain constrained. Analysis of the interface residues (Figure 4B and Supplemental Figure 10A) revealed a pattern of core conservation and peripheral diversification. Specifically, Ser53 and Pro62 are highly conserved across both monocots and dicots (Supplemental Figure 10B), likely providing critical functional and structural integrity—Ser53 for specific hydrogen bonding and Pro62 for maintaining the characteristic peptide turn. Conversely, high variability was observed at peripheral positions, reflecting lineage-specific remodeling where monocots display distinct residue compositions compared to the canonical dicot interface. This divergence is exemplified by Asn56 and Ile61, which exhibit the most significant lineage-specific differences between dicots and monocots (Supplemental Figure 10C and 10D).

To quantify how these lineage-specific changes manifest at the interface level, we computed residue-pair sequence entropies across all receptor–ligand contacts identified in the HSL3–CTNIP4 complex (Supplementary Table 2). These interactions were categorized based on the entropy of both partners: pairs exhibiting low entropy in both the receptor and ligand were classified as co-conserved, whereas those with high entropy were termed co-diversifying (Figure 4C and Supplemental Figure 11). To investigate the structural basis of this resilience, the HSL3–CTNIP4 complex from the monocot *Ananas comosus* was modeled using AlphaFold3 (Figure 4D). Despite low ligand sequence identity, the prediction yielded high confidence scores (ipTM = 0.83, pTM = 0.92), indicating a preserved binding mode. Structural analysis revealed that the core interaction remains intact: the invariant receptor aspartate (Asp288 in AtHSL3, Asp302 in AcHSL3) engages a hydrophilic serine (Ser140) in *A. comosus*, functionally substituting for the canonical asparagine (Asn56). This functional conservation maintains physicochemical compatibility despite sequence variation, thereby enabling HSL3 to retain specific ligand recognition amidst extensive lineage-specific remodeling.

Strikingly, a structure-based DALI search using the HSL3 ectodomain (Supplemental Figure 12) revealed that HSL3 clusters most closely with other peptide-sensing LRR-RKs such as HAESA, MIK2, FLS2, and ERL1. These receptors share a conserved LRR solenoid architecture but differ in overall curvature and ectodomain tilt—features known to sculpt ligand-binding surfaces (Lin et al., 2017; Santiago *et al*., 2016; Sun et al., 2013; Wu *et al*., 2024). This structural diversification across LRR-RKs mirrors the sequence-level patterns observed in HSL3–CTNIP4 evolution, wherein a conserved scaffold is retained while peripheral interface elements diversify to accommodate distinct peptide ligands. The lower similarity of HSL3 to animal LRR proteins further highlights that plant LRR-RKs have undergone plant-specific architectural shifts, likely enabling their expansion into a wide repertoire of peptide-mediated signaling pathways.

Together, these analyses position the HSL3–CTNIP4 module as a representative example of how plant LRR-RKs evolve: by preserving a deeply conserved structural and electrostatic core, while diversifying peripheral contact regions and ligand motifs to tune receptor–ligand compatibility across lineages. This integrated evolutionary framework explains both the robust preservation of HSL3 and the heterogeneous retention of CTNIP4 peptides across plant taxa and provides mechanistic insight into how peptide recognition diversified within the LRR-RK superfamily.

## Discussion

The cryo-EM structures presented here establish a mechanistic framework for how HSL3 senses the stress-induced CTNIP peptides through a recognition mode that diverges substantially from previously characterized LRR–RK systems. While our structural data specifically capture HSL3 in complex with a CTNIP4-derived core fragment, the high conservation of the CTNIP motif and cysteine-stabilized topology suggests that this recognition mode is broadly representative of CTNIP family peptides. Unlike receptors that engage extended and largely linear ligands such as IDA or CLE peptides (Fletcher, 2020; Hohmann *et al*., 2017; Santiago *et al*., 2016), HSL3 recognizes a compact, disulfide-constrained cyclic peptide whose topology is largely pre-organized prior to receptor engagement. This architectural feature minimizes the entropic cost of binding and positions the CTNIP motif in a conformation that is immediately compatible with HSL3’s concave groove, offering an explanation for the relatively fast association kinetics observed in our biochemical assays.

The HSL3 ectodomain adopts a rigidified solenoid architecture, with only a shallow local adjustment around the C-terminal LRRs upon ligand binding. This limited movement suggests that the receptor is preconfigured for rapid ligand capture, a property consistent with the transient and stress-responsive nature of CTNIP accumulation (Liu *et al*., 2022; Rhodes *et al*., 2022b). The electrostatic environment of the inner groove plays a crucial role in this process: the linear Glu357–Arg358–Glu360 cluster forms a guiding surface that positions the positively charged N-terminal flank of CTNIP4 for correct docking. Mutational disruption of this cluster compromises both binding geometry and downstream ROS responses, indicating that electrostatic steering is an essential determinant of ligand orientation and receptor activation. This two-step process—electrostatic guidance followed by motif-anchored stabilization—distinguishes CTNIP perception from other LRR–RK systems that rely more heavily on extensive sequence complementarity along the ligand backbone.

A prominent insight emerging from this study is that glycosylation participates directly in the recognition of cyclic CTNIP peptides by HSL3. Previous work on MIK2–SCOOP complexes has shown that receptor N-glycans can be indispensable for ligand engagement, where they stabilize the receptor surface and shape the architecture of the peptide-binding platform (Jia *et al*., 2024; Wu *et al*., 2024). Our structure reveals a related yet distinct configuration: the Asn449-linked glycan of HSL3 extends toward the ligand and forms a transient contact with Arg63 of CTNIP4, thereby influencing peptide positioning rather than solely reinforcing the receptor scaffold. This mode of glycan participation represents a complementary solution within the LRR-RK family—one in which glycosylation can function not only as a structural buttress, as in MIK2 or FLS2, but also as an active contributor to the microenvironment that modulates ligand orientation and local dynamics (de Oliveira et al., 2016; Haweker *et al*., 2010). The conserved presence and strategic trajectory of this glycan suggest that plant receptors have diversified glycan usage to refine specificity and responsiveness, highlighting glycosylation as a more versatile and nuanced determinant of peptide perception than previously appreciated.

Building on these structural observations, our truncation studies further delineate the minimal features required for productive recognition. While the conserved CTNIP motif forms the rigid core of interaction, the N-terminal residues enhance association rates through electrostatic complementarity and contribute to signaling amplitude in planta. Disruption of the Cys58–Cys68 disulfide bond leads to increased flexibility around the motif and reduced activity, reinforcing the idea that conformational pre-organization is a prerequisite for high-fidelity binding. Taken together, these data reveal that HSL3 achieves specificity not through an extended recognition interface but through the cooperative integration of peptide topology, electrostatic steering, and glycan proximity.

The structural attributes of HSL3 contrast with the architecture of its closest homologs, such as HAESA, which bind linear developmental peptides and undergo more pronounced rearrangements during ligand engagement (Santiago *et al*., 2016). This supports the notion that HSL3 has diverged within the LRR-RK family to accommodate compact, stress-induced peptides rather than developmental cues. Such divergence aligns with the transcriptional behavior of CTNIPs, which are rapidly and transiently induced by pathogen-associated stresses and are also responsive to abiotic stress cues (Bjornson *et al*., 2021; Li *et al*., 2014; Liu *et al*., 2022; Rhodes *et al*., 2022b). The combination of a rigid receptor, a cyclic and pre-organized ligand, and a glycan-tuned pocket may therefore constitute a receptor design optimized for fast on/off signaling cycles, ensuring sensitivity without prolonged activation.

Our evolutionary analysis reveals striking divergence in the HSL3–CTNIPs interface between *Arabidopsis* and the CAM plant *Ananas comosus* (pineapple), despite this module’s critical role in stomatal regulation. While the conserved electrostatic core anchors (D288 and E360 in AtHSL3) are maintained, peripheral interface residues show remarkable lineage-specific remodeling. This divergence is particularly intriguing given that stomatal control—a primary output of HSL3–CTNIPs signaling—has undergone dramatic evolutionary reprogramming in CAM plants. CAM photosynthesis inverts typical stomatal rhythms, opening stomata at night to minimize water loss while fixing CO₂. The extensive sequence divergence we observe at the HSL3–CTNIPs interface in *A. comosus*, including substitutions at positions critical for ligand coordination in Arabidopsis, may reflect adaptive tuning of this signaling module to accommodate the unique temporal dynamics of CAM stomatal regulation. Our AlphaFold3 modeling suggests that despite low sequence identity, the pineapple HSL3–CTNIPs complex maintains a conserved binding mode through compensatory substitutions—for instance, a hydrophilic serine (S140) in A. comosus CTNIP functionally replaces the canonical asparagine at position 56. This evolutionary plasticity highlights how peptide–receptor interfaces can diverge substantially while preserving core signaling functions, potentially enabling fine-tuning of stomatal responses to match distinct physiological strategies across plant lineages.

A recent X-ray crystallographic study by Jiménez-Sandoval et al. reported an HSL3–CTNIP4 complex in which an extended CTNIP4 segment (residues 46–70) was shown to be critical for high-affinity binding (Jimenez Sandoval et al., 2025). Although we did not directly test this longer N-terminal stretch in our truncation series, these observations raise the possibility that residues upstream of position 48 may further refine ligand orientation or contribute to receptor engagement in specific physiological contexts. In this regard, X-ray crystal structure of HSL3 in complex with CTNIP4^46–70^, independently supports the importance of extended N-terminal residues, while our truncation-based mapping uniquely defines the minimal core motif required for productive recognition, highlighting complementary but distinct insights into CTNIP engagement. However, a critical distinction emerges between their crystallographic approach and our cryo-EM analysis: while X-ray crystallography necessitated partial deglycosylation using Endo H to remove non-core glycans for crystal formation, our cryo-EM structure preserves the native glycosylation state of HSL3. This technical difference proved consequential, as we identified a functionally important glycan–peptide interaction at Asn449, where the NAG559 moiety directly contacts CTNIP4 Arg63 through hydrogen bonding. This glycan-mediated contact, which would be absent in deglycosylated crystal structures, contributes to the stabilization of the peptide–receptor interface and may influence the kinetics of ligand binding and release. The preservation of native glycosylation in our cryo-EM analysis thus reveals an additional layer of molecular recognition that extends beyond the protein–peptide interface. This finding underscores the importance of studying receptor–ligand complexes in their native glycosylated state and suggests that receptor glycosylation actively participates in peptide recognition rather than serving merely as a passive structural element. Future studies examining the role of specific glycoforms at Asn449 and other glycosylation sites could provide further insights into how post-translational modifications fine-tune peptide hormone signaling in plants.

Altogether, our findings uncover a distinct structural and mechanistic logic underlying CTNIP perception. HSL3 integrates a topology-constrained peptide, an acidic steering groove, and an active glycan contact to achieve selective yet reversible ligand recognition. This expands the conceptual landscape of peptide–receptor interactions in plants and provides a framework for engineering synthetic ligands or modified receptors with tailored specificities (Li et al., 2025; Zhang et al., 2025). Further examination of co-receptor recruitment and the contribution of glycosylation to higher-order complex assembly will help clarify how this unique recognition module interfaces with downstream signaling machinery.

## Methods

### Plasmid construction

The extracellular domain of HSL3 (AT5G25930, residues 23–662) was generated using *A. thaliana* Col-0 cDNA and was subcloned into a modified pFastBac vector that includes a GP67 secretion signal sequence upstream of a 10XHis tag and the FLAG tag for protein expression in ExpiSf9 cells. All recombination primers were commercially synthesized and purchased from Macrogen (Seoul, Korea). The primers’ sequences are as follows; **pFB-F**: GGATTATTCATACCGTCCCA (20-mer), **pFB-R**: CAAATGTGGTATGGCTGATT (20-mer). The coding sequence of *HSL3* was amplified from *Arabidopsis thaliana* cDNA using gene-specific primers containing overlapping sequences compatible with NcoI-linearized *pCAMBIA1390–35S::Myc*. The PCR fragment was then recombined into the NcoI-digested vector using the In-Fusion HD Cloning Kit (Takara Bio) according to the manufacturer’s instructions. To generate the HSL3^TPM^ variant, point mutations were introduced into *HSL3* by designing mutagenic primers and amplifying two partially overlapping fragments by PCR. These two fragments were fused by overlap-extension PCR to produce the full-length mutated *HSL3^TPM^* sequence, which was subsequently recombined into the NcoI-linearized *pCAMBIA1390–35S::Myc* vector using the In-Fusion cloning system. All resulting constructs were verified by Sanger sequencing.

### Synthetic peptides

All peptides employed in the study were synthesized and procured from GenScript. The peptide sequence is shown below. All peptides were formulated fresh just before preparation of samples and SPR assay. All synthesized peptide sequences are as follows; CTNIP4^48–70^: GPVRGSGRNGCTNIPRGSGRCHN, CTNIP4^51–70^: RGSGRNGCTNIPRGSGRCHN, CTNIP4^54–70^: GRNGCTNIPRGSGRCHN

### Protein expression and purification

The recombinant extracellular domain of HSL3 with a 10XHis tag and flag tag was expressed by Bac-to-Bac baculovirus expression system. Cells (2.0 × 10^6 cells/mL) were infected with the recombinant baculovirus at 20 mL/L and incubated at (22°C). Sixty-eight hours post infection, infected cell culture supernatants were harvested by centrifugation at 1,000 g for 30 min. This supernatant was subjected to Affinity Purification on a HisTrap™ HP 5 mL column (Cytiva, USA) and eluted with the buffer of 50 mM Tris (pH 8.0), 400 mM NaCl, and 500 mM imidazole. The eluant was further desalted using a HiPrep 26/10 desalting column (Cytiva, USA), with a buffer made of 50 mM Tris (pH 8.0). The desalted eluate was concentrated and injected onto a Superdex 200 10/300 GL column (Cytiva, USA) in size exclusion chromatography with a buffer of 1X PBS (pH 7.4). The peak fractions were collected and resolved by SDS-PAGE, protein bands being revealed with Coomassie brilliant blue staining.

### Cryo-EM sample preparation and data acquisition

The concentrate from size exclusion chromatography was concentrated to 1 mM and then diluted to the desired concentration for protein sample preparation using an Amicon® Ultra centrifugal filter (Millipore, USA) with a 30-kDa pore size. HSL3 and CTNIP4^48–70^ were mixed in a molar ratio of 1:3 for the protein-peptide complex preparation. For sample preparation, 3 µL of the HSL3 monomer (0.485 mg/mL) solution was injected on a previously glow-discharged negatively charged holey carbon grid (Quantifoil Au 1.2/1.3, 200 mesh) or the HsL3 – CTNIP4^48–70^ complex (0.630 mg/mL) was injected onto holeycarbon grids (Quantifoil Au 1.2/1.3,300 mesh); both grids had been glow discharged with a negative charge (15 mA) for 60 s using PELCO easiGlow™ glow discharge cleaning system (Ted Pella, USA). Blotted grids (2 s) were plunged into liquid ethane, using a two-sided blotting mode in a Vitrobot Mark IV Thermo Fisher Scientific operated at 4 °C and 100% humidity. All cryo-EM data were collected on a 300-kV Titan Krios G4 Transmission Electron Microscope (Thermo Fisher Scientific, USA). The micrographs were acquired on a Falcon III direct electron detector (Thermo Fisher Scientific, USA) with a nominal magnification of 130,000× (final pixel size of 0.648 Å per pixel). Defocus ranged from -0.7 to -1.8 µm.Raw data were further processed in cryoSPARC version 4.5.3 (Punjani et al., 2017).

### Image processing and 3D reconstruction

In cryoSPARC (Punjani *et al*., 2017), 21,554 micrographs for HSL3 monomer and 8,404 for the complex of HSL3 – CTNIP4^48–70^ were imported independently. The imported movies were motion corrected and CTF-estimated in full frame, and poor quality micrographs were manually excluded. The downstream workflows for the HSL3 monomer and HSL3 – CTNIP4^48–70^ complex are described in a supplemental Figure 2 and 3. A total of 15,508 and 5,858 movies were collected for the HSL3 monomer and HSL3 – CTNIP4^48–70^ complex, respectively. Batch 2D classification outputted 2,900,290 and 961,291 particles for monomeric HSL3 and the HSL3 – CTNIP4^48–70^ complex respectively which were used for ab initio reconstruction consisting of two classes each. For the HSL3 monomer, a total of 2,101,488 particles were used for the initial 3D classification and ultimately 1,740,510 particles were selected for final 3D refinement. For the HSL3 – CTNIP4^48–70^ complex, first 3D classification repeated twice (731,410 particles) followed by final selection of 350,460 particles for 3D refinement. After post-processing, the final resolution of the 3D reconstructions for HSL3 monomer and HSL3 - CTNIP4^48–70^ complex were 2.63 Å and 2.59 Å, respectively.

### Model building and refinement

An initial model of *Arabidopsis* HSL3 was generated using the AlphaFold-predicted structure as a starting template. The model was manually fitted into the cryo-EM density map using ModelAngelo (Jamali et al., 2024) implemented within UCSF ChimeraX (version 1.6) (Pettersen et al., 2021) to guide the overall placement of the receptor domain. Subsequent manual model building and adjustment were performed in Coot (version 0.9.8.1) (Emsley et al., 2010) by iteratively refining side-chain conformations and correcting backbone traces according to the density. Real-space refinement was carried out using Phenix.real space refine (version 1.20) (Liebschner et al., 2019), employing secondary structure and Ramachandran restraints throughout the refinement cycles. Glycosylation at each Asn residue was modeled in Coot using the built-in glycosylation module, with *N*-acetylglucosamine (NAG) residues positioned such that the Asn ND2 atom was within ∼1.5 Å of the NAG C1 atom, consistent with the density and expected geometry. Covalent linkages between the Asn side chain and the attached NAG moieties were generated using the Make Covalent Link (AceDRG) (Long et al., 2017) utility in CCP4i2 (Potterton et al., 2018) to ensure proper bonding restraints prior to final refinement. The final refined model was validated using the standard Phenix and CCP-EM validation tools, and model statistics, including map-to-model correlation and geometry metrics, are summarized in Supplementary Table 1.

### Surface Plasmon Resonance (SPR) assay

HSL3 binding kinetics and affinities to CTNIP4 variants were measured on a Biacore T200 (GE Healthcare, United States) instrument with CM5 chips (Cytiva, USA) at 25°C. HSL3 proteins were exchanged into 10 mM NaAc pH 5.0 and the peptides were dissolved in HBS-EP+ used as running buffer [10 mM Hepes, 150 mM NaCl, 3mM EDTA and added with 0.05% (v/v) Surfactant P20 pH7.5] (Cytiva,. USA). 3,500 response units of HSL3 proteins were captured on the CM5 chip, using a blank channel as negative control. The peptides were diluted to the indicated concentrations and loaded at a flow rate of 30 µL min^-1 for 120 seconds, and then dissociated for 600 seconds. Glycine (pH 1.5) was then injected for 30 s following dissociation to clear any residual peptides from the chip surface. Binding kinetics was determined by Biacore T200 Evaluation Software version 3.2 using 1:1 Langmuir binding model .

### Global N-Glycan Profiling

N-glycans were released by sequential digestion with PNGase F and then PNGase A, and purified on a porous graphitized carbon (PGC) cartridge (100 mg, 1 mL tube). The purified glycans were combined (1:1, v/v) with 2,5-dihydroxybenzoic acid (5 mg/100 μL in water-50% acetonitrile) and spotted on a MALDI plate made of stainless steel. The vacuum dried samples were analyzed using a MALDI-8030 TOF MS (Shimadzu, Japan) in the m/z 500 – 3000 range.

### Site-Specific N-Glycan Analysis

Purified protein (100 μg) was reduced with 2 μL of 550 mM dithiothreitol in 50 mM ammonium bicarbonate buffer (1:1, *v/v*) at 60 °C for 50 min and alkylated with 4 μL of 450 mM iodoacetamide in the dark at room temperature for 45 min. After adjusting the pH to 7 – 8, samples were digested with trypsin and Glu-C (enzyme to substrate ratios of 1:100 and 1:50, *w/w*) at 37 °C for 3 h. Glycopeptides were enriched using an Oasis MAX cartridge (30 mg, 1 mL tube) then completely dried in vacuum. Purified N-glycopeptides were reconstituted in 0.1% formic acid and analyzed using a ZenoTOF 7600 MS (SCIEX, USA) coupled to an ACQUITY UPLC M-Class system (Waters, USA). Peptides were trapped on an Acclaim PepMap100 C18 column (75 μm × 20 mm, 3 μm, 100 Å) and separated on a BEH C18 column (75 μm × 150 mm, 1.7 μm, 130 Å) at 300 nL/min. Mobile phases were 0.1% formic acid in water (A) and 0.1% formic acid in acetonitrile (B) with a 130 min gradient from 4% to 96% B. MS spectra were acquired over *m/z* 400 – 3,000 (0.2 s accumulation), and MS/MS spectra were collected over *m/z* 150 – 3,000 using collision energy of 20 ± 5 and 0.05 s accumulation. Precursor ions were isolated with a 2.0 *m/z* window. Raw data were processed using Byos software (Protein Metrics) against the UniProt database. Glycan compositions obtained from MALDI–TOF MS were used to generate a custom glycan library, and all spectra were manually verified for confident glycopeptide assignment.

### Plant material and growth conditions

*Arabidopsis thaliana* accession Columbia-0 (Col-0) was used as the wild-type control in all experiments performed. The T-DNA insertion mutant *hsl3-1* (SALK_207895C) was described previously (Liu et al., 2020) and was obtained from the *Arabidopsis* Biological Resource Center (ABRC). *Arabidopsis* plants for ROS burst assays, were grown individually in pots filled with soil in a controlled growth room at 22℃, 70% relative humidity, with a 16 h light/8 h dark photoperiod and a light intensity of 120-140 µmol m^-2^ s^-1^. Plants were used at four-to five-week-old. Seeds used for analyses of growth inhibition and MAPK activation were surface-sterilized by sequential washing in 70% ethanol and 100% ethanol, followed by three rinses with sterile distilled water, and then stratified in the dark at 4 ℃ for 1 day. Sterilized seeds were sown on half-strength Murashige and Skoog (1/2 MS) medium (1% sucrose, 0.8% plant agar, pH 5.7), and were germinated and grown for 1 week under the same temperature, humidity, light intensity, and photoperiod as soil-grown plants. *Nicotiana benthamiana* plants used for *Agrobacterium*-mediated transient expression and ROS assays were grown on soil under the same growth conditions described above.

### Transient expression in *N. benthamiana*

All binary plasmids were transformed into *Agrobacterium tumefaciens* strains C58C1 for transient expression in *N. benthamiana*. *Agrobacterium* cultures were grown overnight at 28 °C in liquid Yeast Extract Peptone (YEP) medium supplemented with appropriate antibiotics, including kanamycin, and rifampicin corresponding to the binary vector backbone. Cells were collected by centrifugation at 3,000 × g for 10 min and resuspended in infiltration buffer consisting of 10 mM MES–KOH (pH 5.7), 10 mM MgSO_4_, and 100 µM acetosyringone. The resuspended cultures were incubated at room temperature for 3 h, after which cell suspensions were adjusted to a final OD₆₀₀ of 0.6 and mixed with *Agrobacterium* strains carrying the receptor construct and the P19 silencing suppressor at a 4:1 ratio. The mixtures were subsequently infiltrated into the first fully expanded leaves of *N. benthamiana* using a 1-mL needleless syringe. Control samples were infiltrated with P19-only *Agrobacterium* suspensions.

### Reactive oxygen species (ROS) assay

For ROS burst assays, leaf discs were taken from 4-week-old *Nicotiana benthamiana* and *Arabidopsis thaliana*. For *N. benthamiana*, AtHSL3 and AtHSL3^TPM^ (E357A/R358A/E360A) were transiently expressed by *Agrobacterium* infiltration, and discs were collected 48 h post-infiltration using a 4-mm diameter biopsy punch (KAI MEDICAL). For *A. thaliana*, leaf discs of the same diameter were excised without prior infiltration. Discs were placed abaxial side up in white 96-well plates (SPL Life Sciences) containing 200 μL of sterile water. During the first hour, the water was replaced twice at 30-min intervals, after which the plates were incubated overnight at 22 °C in the dark. After incubation, water was removed, and reactions were initiated by adding assay solution containing luminol L-012 (100 µM; FUJIFILM Wako Chemicals USA, Cat# NC0733364), horseradish peroxidase (20 µg mL⁻¹; Sigma-Aldrich, Cat# 516531-5KU), and the indicated elicitor peptide (1 µM CTNIP4^48–70^, CTNIP4^51–70^, or CTNIP4^54–70^) to a final volume of 100 µL per well. Chemiluminescence was recorded on an Infinite® M Plex multimode microplate reader (Tecan) with 1-min intervals (integration time 300 ms) for 60 min at room temperature (Lee et al., 2024).

### Seedling growth inhibition (SGI) assay

One-week-old seedlings grown on vertical 1/2 MS plates were transferred into 48-well plates (Costar, Corning), placing one seedling in each well. Each well contained 1 mL liquid ½ MS medium supplemented with 1% (w/v) sucrose, and the plates were incubated for 2 days at 22 °C under a 16 h light/8 h dark photoperiod. The medium was then replaced with fresh 1/2 MS containing 1% sucrose, supplemented with one of the three N-terminally truncated CTNIP4 variants (CTNIP4^48-70^, CTNIP4^51-70^, CTNIP4^54-70^) at a final concentration of 5 μM, or with unsupplemented medium for mock controls. One seedling was placed in each well, and a total 8 or 24 seedlings was used for each treatment. The plates were returned to the same growth conditions and incubated for 1 week, after which seedlings were gently dried on paper towels and fresh weigh was measured (Lee *et al*., 2024).

### Sequence curation

Orthologous HSL3 receptors and CTNIP peptides were curated from the resources of Rhodes et al. (Rhodes et al., 2022a) and cross-checked against public annotations when available. HSL3 ectodomains (ECDs) were extracted using *Arabidopsis thaliana* HSL3 as the reference for residue numbering. CTNIPs were trimmed to the AtCTNIP4 motif window (positions 37–70) to harmonize positional indexing. Species with incomplete ECDs, clear frame errors, or ambiguous peptide segments were excluded.

### Multiple sequence alignments

Protein multiple sequence alignments (MSAs) for HSL3 ECDs and CTNIPs were generated with MUSCLE (v5; default parameters) (Edgar, 2004) and visually inspected at interface-proximal sites. Alignments were anchored to the *Arabidopsis thaliana* references (AtHSL3 or AtCTNIP4), and gap characters at the reference position were treated as missing values.

### Phylogenetic analysis and clade annotation

Guide trees were inferred in MEGA (v12) (Kumar et al., 2024) using maximum likelihood under MEGA defaults with 500 bootstrap replicates. Trees were used for visualization and clade labeling only (*Poaceae*, monocots, dicots) and were not used for hypothesis testing. Clade brackets were overlaid on the dendrograms in the plotted heatmaps.

### Reference-based identity scoring and heatmaps

For each aligned position in AtHSL3 or AtCTNIP4 we scored each homolog as 1 when the residue matched the Arabidopsis reference and 0 when it did not; gaps were set to NA. For species with multiple homologs we reported the mean of these 0/1 scores, giving a species-level fraction identical at that site (0–1). Heatmaps plot these fractions by species and site. The analysis covered ten HSL3 ectodomain sites (Q120, Q169, E218, E219, D241, D288, N312, E334, R358, E360) and the CTNIP motif 37–70, including the nine interface sites G48, R51, S53, R55, N56, T59, N60, I61 and P62.

### Sequence logo generation

Amino-acid frequencies were computed from the MSAs after excluding gaps at the plotted columns. Sequence logos were generated in R (ggseqlogo) (Wagih, 2017) with residues colored by biochemical class (acidic, basic, polar uncharged, hydrophobic). For clade-specific panels, the same procedure was applied to monocot-only and dicot-only.

### Shannon entropy and paired-site analysis

Per-position conservation was quantified as Shannon entropy (Shannon, 1948). For structure-defined interacting pairs (HSL3-site ↔ CTNIP-site), receptor and ligand entropies were plotted as mirrored bars on a common scale. Pairs with both entropies ≥ 0.50 were annotated as co-variable; pairs with both < 0.50 were annotated as co-conserved.

### Structure-guided mapping of interacting residues

Interacting residues in the AtHSL3–CTNIP4 complex were defined on our refined atomic model using a heavy-atom distance cutoff of ≤ 4.0 Å. The resulting AtHSL3 and AtCTNIP4 positions were then transferred to orthologs by column mapping through the MUSCLE MSAs. For cross-species models (e.g., *Ananas comosus*), the corresponding residues were identified by alignment to these structure-defined interface positions. AlphaFold3 predictions were used to visualize the interface (Abramson et al., 2024). UCSF ChimeraX (v1.8) was used only for molecular rendering and figure preparation (Pettersen *et al*., 2021).

### Structural similarity analysis and dendrogram construction

The ectodomain structure of *Arabidopsis thaliana* HSL3 (HSL3 ALL-IN model) was submitted to the DALI server (Holm et al., 2023) to identify structurally related proteins in the Protein Data Bank (PDB). Pairwise Z-scores were retrieved and ranked, and the top 100 highest-scoring structures were selected for comparative analysis. A pairwise Z-score matrix was converted to a structural distance matrix using the transformation distance = (max Z-score − Z-score). Hierarchical clustering was performed using UPGMA (average linkage), and the resulting tree was exported in Newick format. The Newick tree and Z-score values were imported into the Interactive Tree of Life (iTOL v6) (Letunic and Bork, 2024). Z-score values were mapped onto branches as a color gradient (green = high similarity; red = low similarity). Proteins were further grouped by similarity rank, and representative structures from each group were manually annotated. The final circular dendrogram visualizes the structural relationship of HSL3 to plant LRR-RKs and other LRR-containing receptors.

## Data availability

Coordinates for the HSL3 apo and HSL3–CTNIP4 complex structure have been deposited under PDB 9XNU and PDB 9XNT. Cryo-EM maps for the HSL3 apo and HSL3–CTNIP4 complex structure are available under EMDB–67060 and EMDB–67059.

## Author contributions

H.J. and H.R. performed gene cloning and protein purification. H.J. and J.-H.Y. carried out cryo-EM structure determination. H.J., Y.K., and H.R. performed SPR experiments. N.S., Y.H.N., and H.J.A. carried out MALDI-MS experiments. H.R., S.C., K.D., and K.A. conducted plant experiments. D.-H.L. and J.H.K. supervised plant experimental design and methodology. H.J., H.R., S.C., J.-H.Y., and H.- S.L. wrote and edited the original draft. B.-K.K. reviewed and edited the manuscript. H.J., H.R., S.C., and J.-H.Y. designed the original draft figures. J.-H.Y. and H.-S.L. refined and finalized the figures.

## Acknowledgments

We thank all members of the IMGN at Kyung Hee University for valuable discussions and technical support. We are grateful to the cryo-EM facility at IBS for assistance with data collection and the plant facility team at IBS for maintaining growth chambers and providing excellent plant care.

