## Supplemental Figure for "Structural basis of cyclic phytocytokine recognition by the HSL3/NUT receptor"

### Supplemental information

**Supplemental Figure 1. Purification and validation of the HSL3 ectodomain.**

**(A)** Size-exclusion chromatography (SEC) profile of purified HSL3 ectodomain monitored at 280 nm, showing a single, monodisperse peak corresponding to the correctly folded protein. **(B)** SDS–PAGE (Coomassie-stained, left) and anti-FLAG immunoblot (right) of the purified protein. Molecular weight markers (kDa) are indicated on the left. Both analyses confirm high purity and the expected molecular weight of glycosylated HSL3.

**Supplemental Figure 2. Cryo-EM data processing flowchart and map quality of apo HSL3.**

**(A)** Flow chart showing the cryo-EM data processing procedure. A representative micrograph (top right) and sets of representative 2D class averages (middle right) are shown. The data were processed with a series of steps for motion correction, contrast transfer function (CTF) parameter estimation, particle picking, 2D classification, *ab initio* reconstruction, homogeneous refinement, heterogeneous refinement, and non-uniform refinement in cryoSPARC. **(B)** Reconstructed 3D map of apo HSL3 shown with local resolution estimate and the final particle number. The density map showed clear features of the LRR domain and terminal caps. **(C)** Gold-standard FSC (GSFSC) curve determined by using the 0.143 criterion represents a global resolution of 2.63 Å. The corrected FSC curve is displayed in purple.

**Supplemental Figure 3. Cryo-EM Data Processing Flowchart and Map Quality of the HSL3–CTNIP4 Complex.**

**(A)** Flowchart depicting the cryo-EM data processing Workflow for the HSL3–CTNIP4 complex. A representative micrograph is shown in the upper right together with sets of representative 2D class averages on the middle right. Data were processed via multiple steps, for motion correction (CTF estimation, particle picking, 2D classification and ab initio reconstruction) homogeneous refinement heterogeneous refinement non-uniform refinement on cryoSPARC. **(B)** Reconstructed 3D map of the HSL3–CTNIP4 complex is shown alongside local resolution estimation and the final particle number used. The density map clearly delineates the overall structure of HSL3 in complex with CTNIP4, displaying well-defined features including the receptor leucine-rich repeat (LRR) domain and bound peptide density along the concave surface. **(C)** The gold-standard Fourier shell correlation (GSFSC) curve, calculated with the 0.143 criterion, is indicative of a global resolution of 2.59 Å. The corrected FSC curve after masking is shown in purple.

**Supplemental Figure 4. Cryo-EM density of CTNIP4 peptide bound to HSL3.**

High-resolution cryo-EM density of the CTNIP4$^{48–70}$ peptide (grey mesh, contoured at a representative threshold) fitted within the HSL3 binding groove. The peptide model is shown as red sticks, with its N- and C-termini labeled. The CTNIP motif region is indicated, highlighting its well-defined density and compact conformation within the receptor pocket. The map clearly resolves side-chain features along the peptide backbone, including the disulfide linkage (Cys58–Cys68) depicted in yellow sticks.

**Supplemental Figure 5. Electrostatic surface potentials of HSL3/CTNIP4 complex.**

**(A)** Electrostatic surface potential maps of the HSL3–CTNIP4 complex shown from multiple orientations. Surfaces are colored according to electrostatic potential (red, negatively charged; blue, positively charged). These views illustrate how the concave face of HSL3 provides an extended acidic landscape that accommodates CTNIP4 upon binding. **(B)** Close-up electrostatic view of the CTNIP4-binding region. The negatively charged region (yellow dashed outline) and the adjoining hydrophobic groove (black dashed outline) are highlighted to illustrate the two principal surface features that guide CTNIP4 docking into the receptor pocket.

**Supplemental Figure 6. Structural comparison between apo and CTNIP4$^{48–70}$ bound HSL3.**

The Cα–Cα residue-wise distance-difference matrix ($\Delta d = d_{(apo)} - d_{(complex)}$) visualizes pairwise conformational changes across the ectodomain. Positive r-DDM values (red) indicate residue pairs that are closer in the complex than in the apo structure, reflecting local contraction upon peptide binding. Negative values (blue) mark residue pairs that are closer in the apo state, indicating regions that expand or shift outward in the complex. Values near zero (white) denote minimal structural change.

The ΔB-factor (Complex – Apo) represents the per-residue difference in average B-factors between the apo and CTNIP4$^{48–70}$ bound HSL3 structures. Negative ΔB values indicate reduced B-factors in the complex, suggesting ligand-induced stabilization or rigidification of these regions. Conversely, positive ΔB values reflect increased B-factors in the complex, consistent with locally enhanced flexibility or partial loosening.

**Supplemental Figure 7. HSL3 and mt HSL3 expression in *Nicotiana benthamiana* leaves.**

Leaf discs (4 mm) transiently expressing AtHSL3 or the triple mutant HSL3(E357A/R358A/E360A) were harvested after the ROS assay; 12 discs per condition were pooled. Proteins were resolved by SDS–PAGE and immunoblotted with an anti-Myc HRP–conjugated antibody (exposure 3 s). Top: anti-

Myc immunoblot showing accumulation of the expressed receptors. Bottom: Ponceau S staining of the membrane as a loading/transfer control. Lanes show independent samples for each condition

**Supplemental Figure 8. Positive ion MALDI–MS spectrum of N-glycans released from the HSL3 ectodomain.**

N-glycans were enzymatically released from the HSL3 ectodomain containing twelve N-glycosylation sites and analyzed by positive ion MALDI–MS. Major glycoforms are labeled using the composition code format (Man_HexNAc_Fuc), and representative structures are depicted following the Symbol Nomenclature for Glycans (SNFG): green circles, mannose (Man); blue squares, N-acetylglucosamine (GlcNAc); red triangles, fucose (Fuc). This spectrum illustrates the overall distribution pattern of high-mannose–type glycans predominantly observed in HSL3.

**Supplemental Figure 9. HSL3 ectodomain conservation and interface-site sequence logos across species and clade.**

**(A)** Conservation of HSL3 residues at the CTNIPs interface (receptor side). Heatmap shows reference-based identity to AtHSL3 for all receptor residues that contact CTNIP in interface map with both bonding (hydrogen bond, salt bridge, or polar contact $\leq$ 4.0 Å) and non-bonded Van der Waals contacts ($\leq$ 4.5 Å). Positions (AtHSL3 numbering): Y214, W216, E218, E219, V244, Y265, F267, A268, D288, V310, F314, E334, R358, E360, G382, V384, Y386, L405, Y429. Identity was scored per homolog (1 if identical to AtHSL3, 0 otherwise; gaps = NA) and averaged per species (0–1). Rows are ordered by a maximum-likelihood guide tree from the HSL3-ECD MSA. Color scale: 0–1; gray, NA/gap. **(B)** HSL3 interface residues in all species (sequence logo). Sequence logo from the HSL3 ectodomain alignment at CTNIP-contact residues (AtHSL3 numbering): Y214, W216, E218, E219, V244, Y265, F267, A268, D288, V310, F314, E334, R358, E360, G382, V384, Y386, L405, Y429. Letter height reflects amino-acid frequency (0–1); colors denote residue class (acidic, basic, hydrophilic, hydrophobic). **(C)** Dicot subset. As in (B), but computed from the dicot HSL3 alignment at the same interface positions (Y214–Y429). Letter height indicates frequency; colors show residue class. **(D)** Monocot subset. As in (B), but for the monocot HSL3 alignment at the same interface positions. Letter height indicates frequency; colors show residue class.

**Supplemental Figure 10. CTNIP4 motif conservation and interface-site sequence logos across species and clades.**

**(A)** Conservation of CTNIP residues at the HSL3 interface (ligand side). Heatmap shows reference-based identity to AtCTNIP4 across positions 37–70. Identity was scored per homolog (1 if identical to AtCTNIP4, 0 otherwise; gaps = NA) and averaged per species (0–1). Black boxes mark interface segments, and underlined x-axis labels denote CTNIP residues that contact HSL3 in interface map with both bonding (hydrogen bond, salt bridge, or polar contact ≤ 4.0 Å) and non-bonded Van der Waals contacts (≤ 4.5 Å). Color scale: 0–1; gray, NA/gap. **(B)** CTNIP interface residues in all species (sequence logo). Sequence logo from the CTNIP alignment at HSL3-contact residues (AtCTNIP4 numbering): R51, G52, S53, R55, N56, N60, I61, P62, R63, H69. Letter height reflects amino-acid frequency (0–1); colors denote residue class (acidic, basic, hydrophilic, hydrophobic). **(C)** Dicot subset. As in (B), but computed from the dicot CTNIP sequences at the same interface positions. Frequencies shown as letter heights; colors by residue class. **(D)** Monocot subset. As in (B), but for the monocot CTNIP sequences at the same interface positions. Frequencies shown as letter heights; colors by residue class.

**Supplemental Figure 11. Sequence entropy of HSL3–CTNIP interface residue pairs (all contacts).**

**(A)** Sequence entropy for receptor–ligand interface pair, including both bonding (hydrogen bond, salt bridge, or polar contact ≤ 4.0 Å) and non-bonded Van der Waals contacts (≤ 4.5 Å). Receptor entropies were computed from the HSL3 ectodomain MSA; ligand entropies from the CTNIP residues 37-70 alignment. Vertical dashed lines mark the median entropy of each set (receptor, ligand). Pairs with both entropy < median are classified as co-conserved (yellow star); pairs with both entropy ≥ median are co-diversifying (red star). Bars are labeled with entropy values; residue names follow AtHSL3/AtCTNIP4 numbering.

**Supplemental Figure 12. Structural similarity of HSL3 relative to plant LRR-RKs and other related receptors**.

A structural dendrogram was generated using DALI pairwise Z-scores with the *Arabidopsis thaliana* HSL3 ectodomain (HSL3 ALL-IN model) as the reference. Because DALI compares only structural similarity, the Z-score heat scale reflects the degree of resemblance to HSL3: green indicates high similarity, red indicates low similarity. Notably, Z-scores above 40 represent structures that are nearly identical in overall fold. Based on this scale, proteins with very high structural similarity to HSL3 are highlighted in purple, those with substantial similarity in green, and those with moderate but notable similarity in blue. Only representative proteins from each group are labeled in both the table and dendrogram for clarity (labels shown in red). The proteins most structurally similar to HSL3 include: *Arabidopsis thaliana* HAESA, *Brassica napus* MIK2, *Arabidopsis thaliana* FLS2, *Arabidopsis thaliana* ERL1, which cluster closely with HSL3 in the dendrogram.

**Supplemental Table 1. Cryo-EM data collection, refinement, and validation statistics.**

**Supplemental Table 2. Comprehensive interaction map of the HSL3–CTNIP4 Binding Interface.**

### Supplementary Figure 1

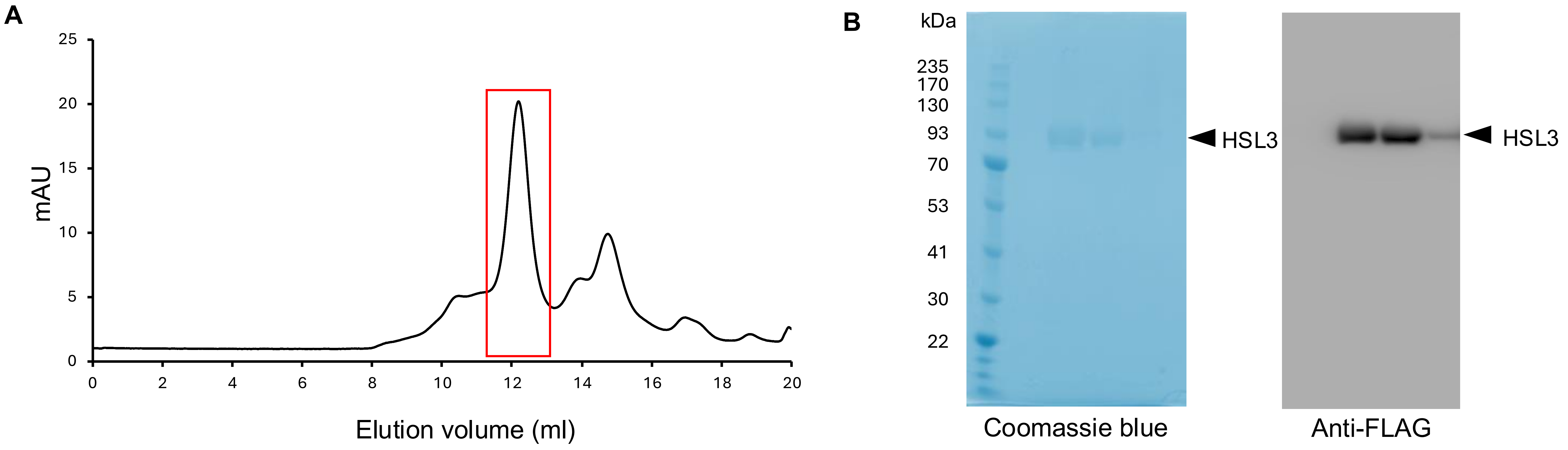

### Supplementary Figure 2

**A**

**21,554 movies** (300kV, SA 2.7 mm, 0.648 apix, 65.6 e/Å^2) ▶

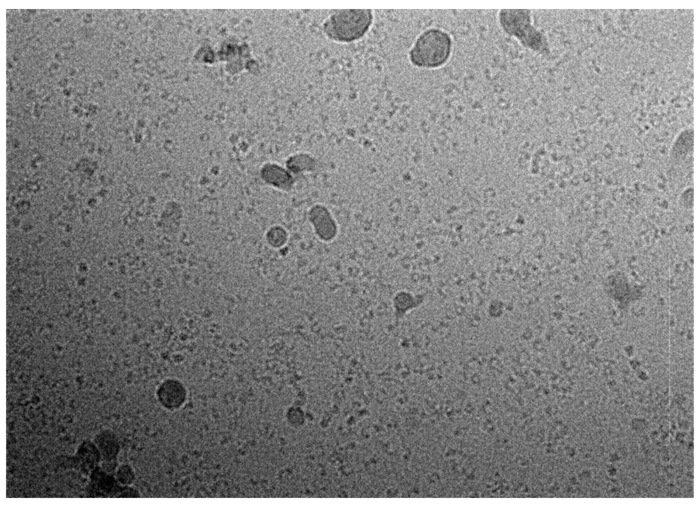

Nominal magnification : X 130,000

↓
Patch motion correction (multi)
↓
Patch CTF estimation (multi)
↓
Curate exposure (**15,508 exposure accepted**)
↓
Blob picking using 15,508 micrographs, 12,913,057 particles → 6,925,859 particles selected
↓
2D classification with 50 classes, selected 26 classes, total 4,318,575 particles ▶

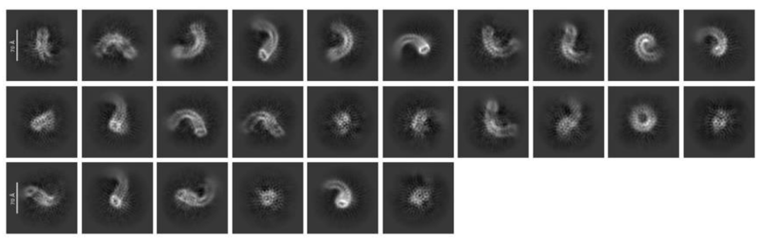

↓
Template picking using 26 classes, 15,508 micrographs; total 12,145,753 particles → 8,038,044 particles selected
↓
2D classification with 50 classes, selected 17 classes, total 3,593,817 particles ▶

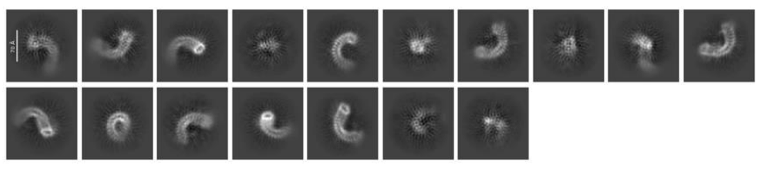

↓
2D classification with 50 classes, selected 35 classes, total **2,900,290 particles** ▶

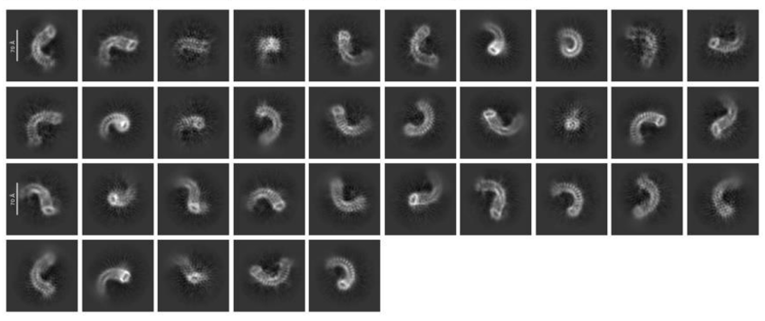

↓
Ab-initio reconstruction with 2 classes, **2,101,499 particles selected**
↓
**Homogeneous refinement**

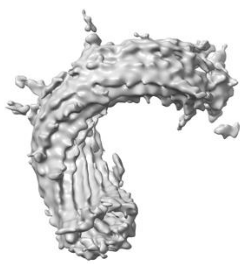

3.51Å

**1st Heterogeneous refinement**

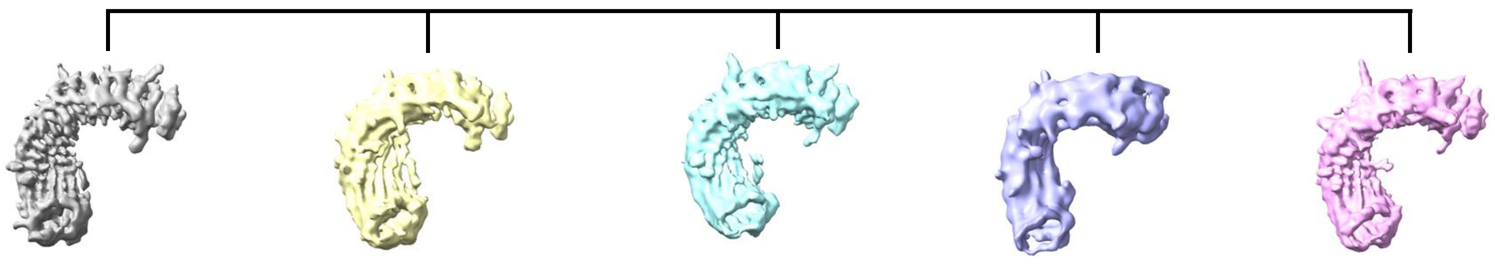

**601,304 particles** | **451,633 particles** | **369,401 particles** | **360,989 particles** | **318,172 particles**

**Homogeneous refinement, Non-uniform refinement**

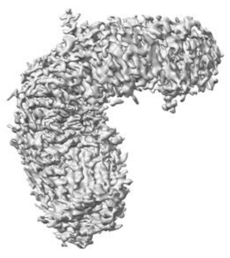

**B**

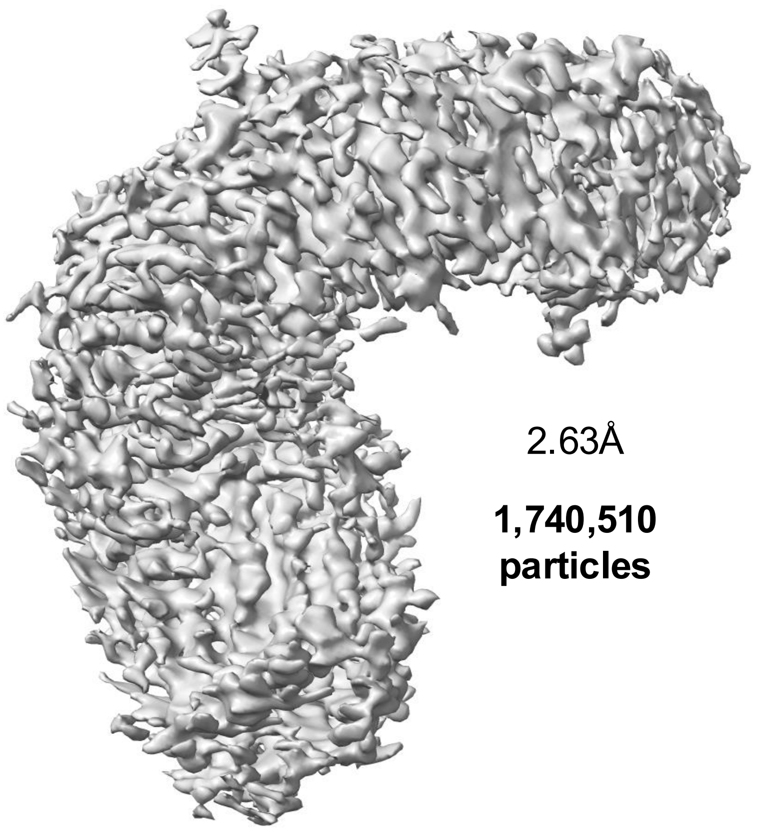

2.63Å

**1,740,510 particles**

**C**

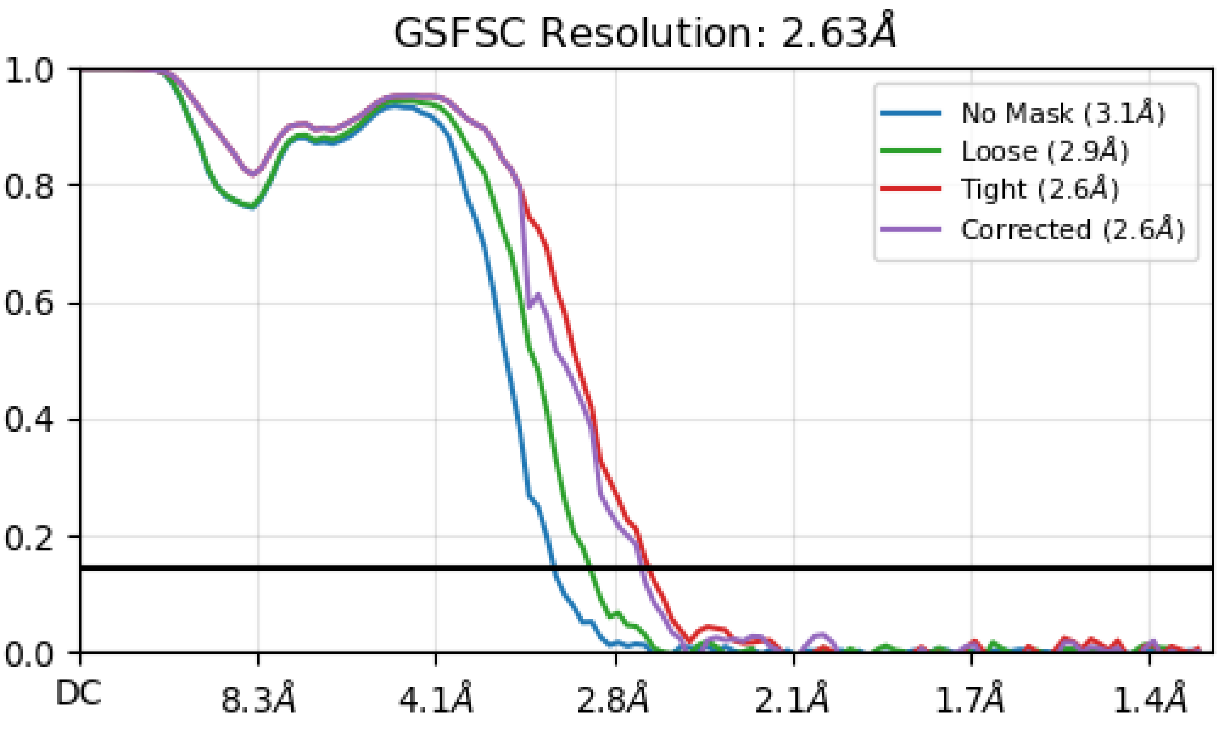

### Supplementary Figure 3

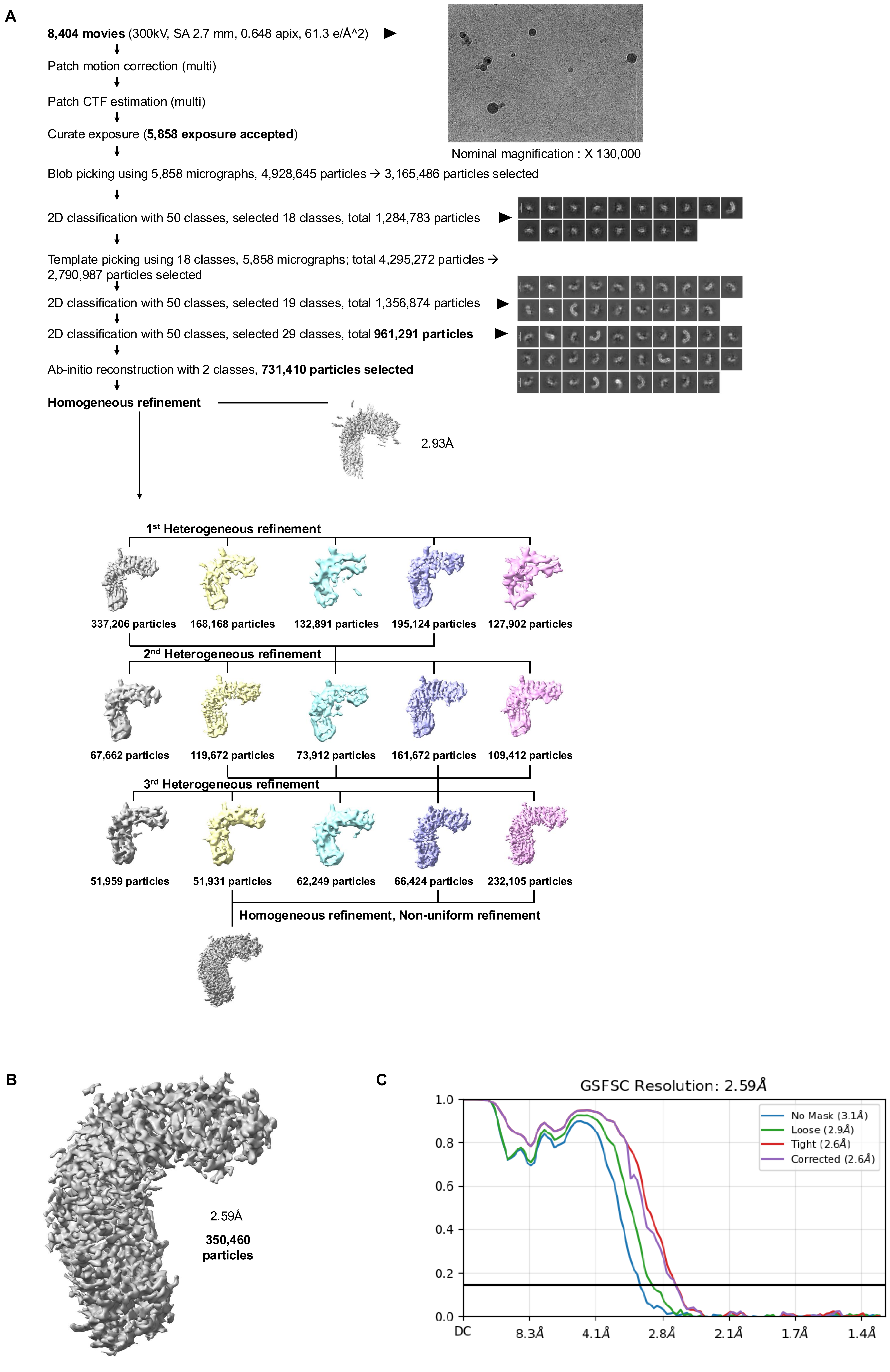

### Supplementary Figure 4

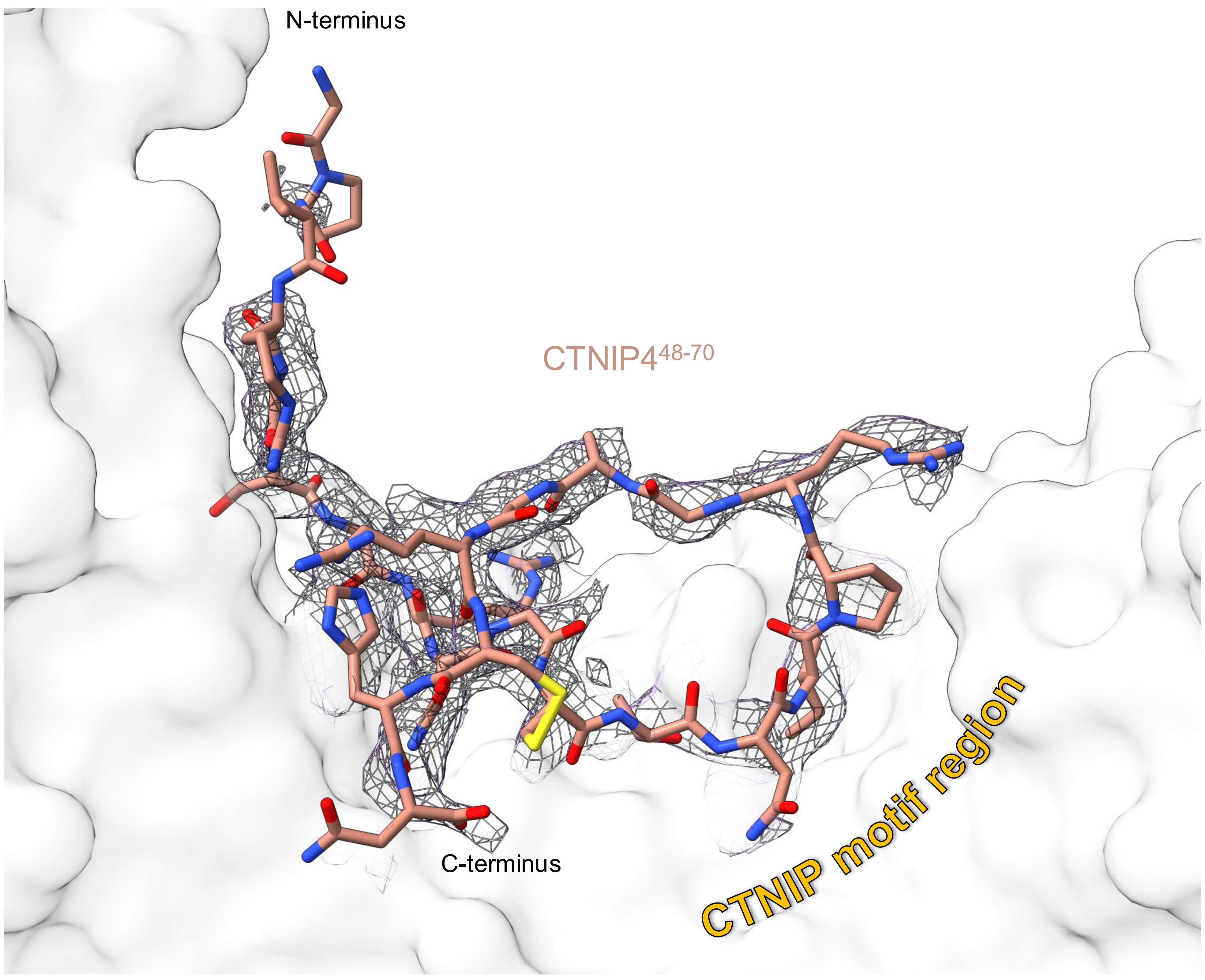

Supplementary Figure 5

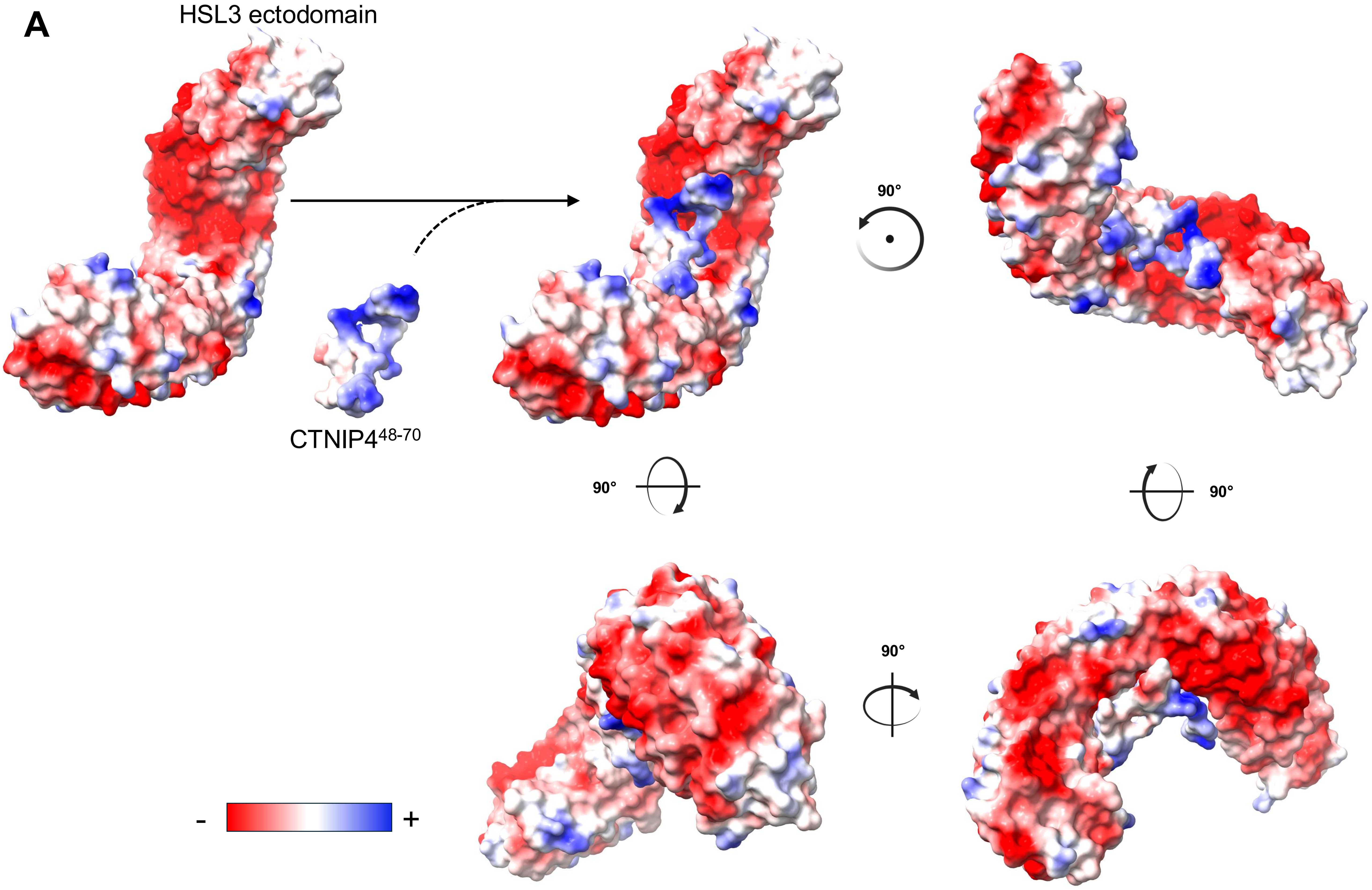


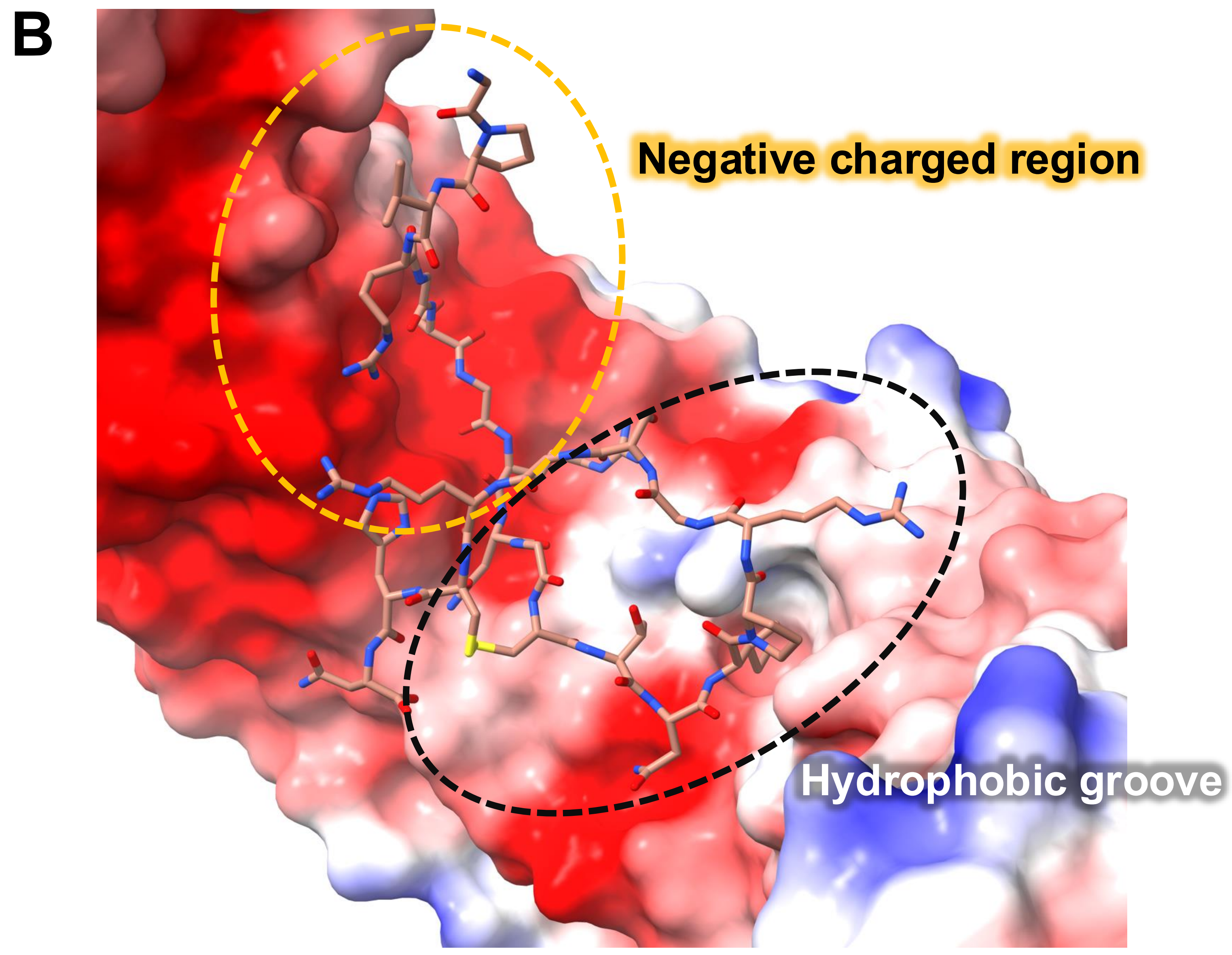

Supplementary Figure 6

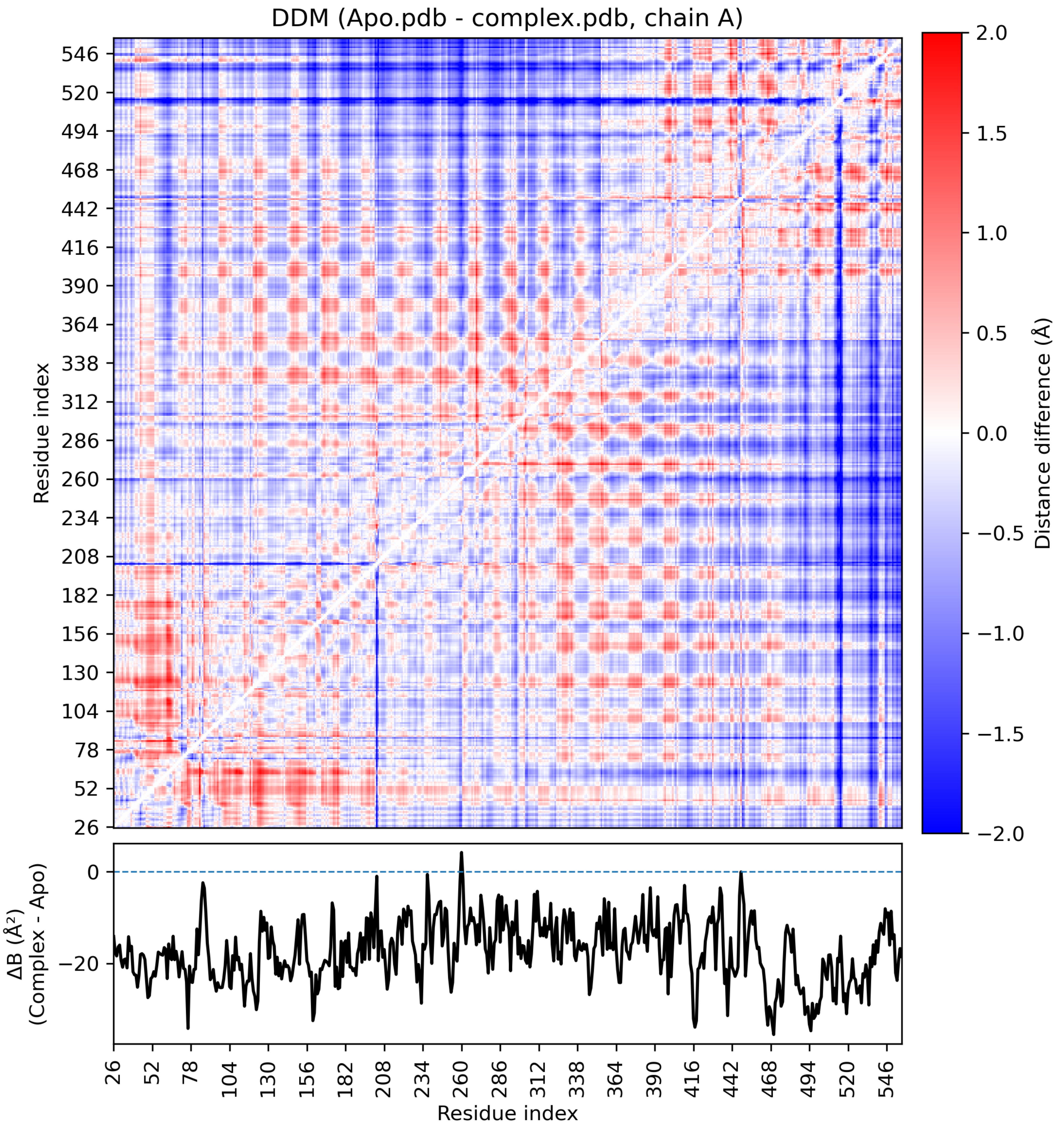

### Supplementary Figure 7

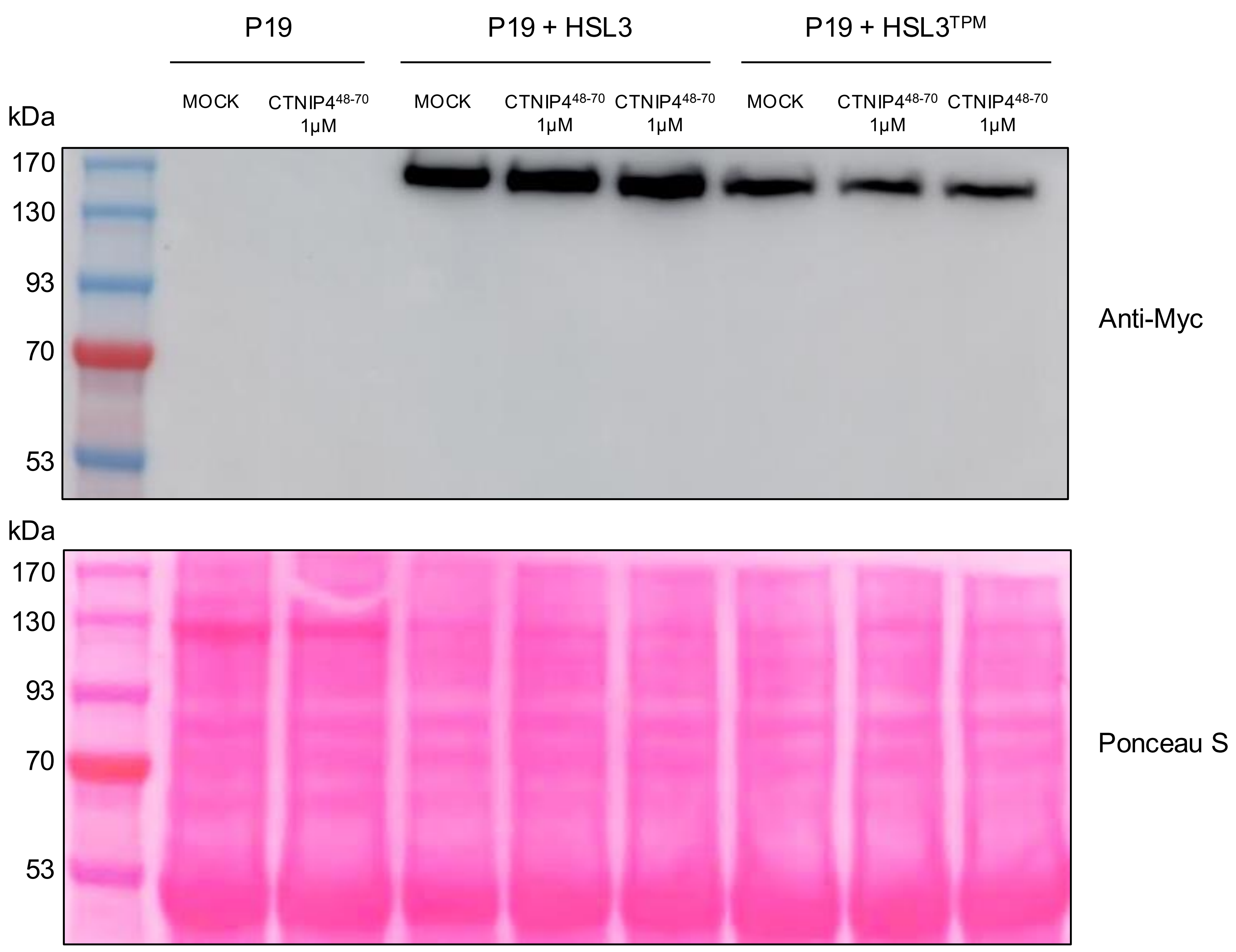

### Supplementary Figure 8

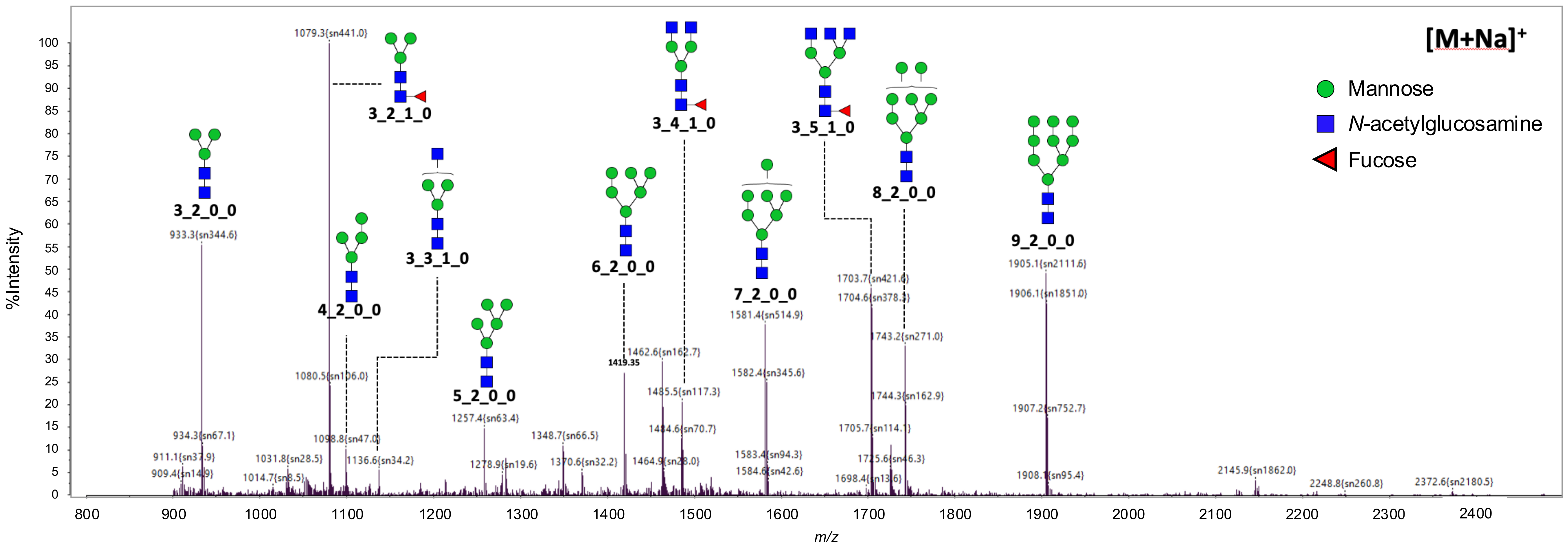

### Supplementary Figure 9

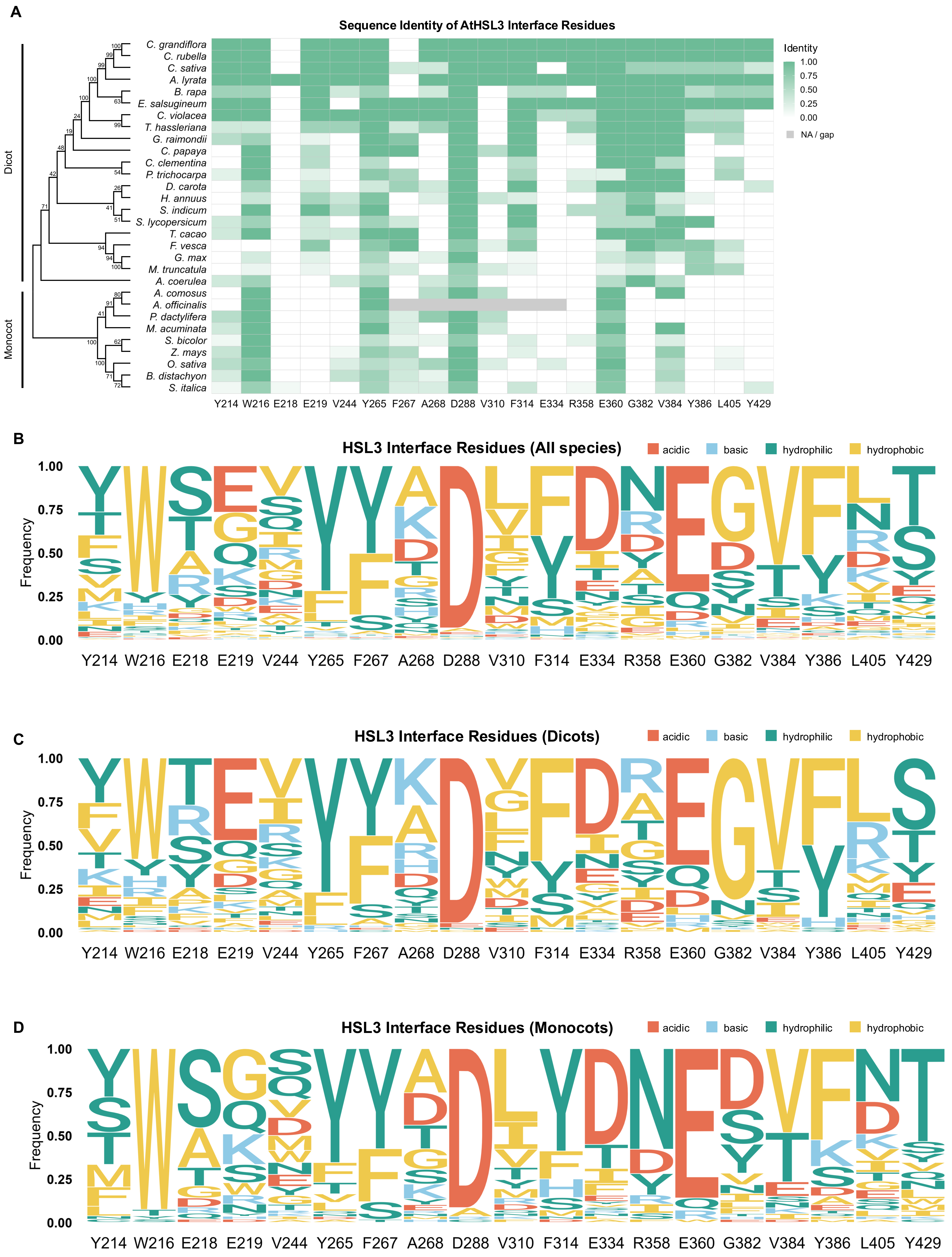

### Supplementary Figure 10

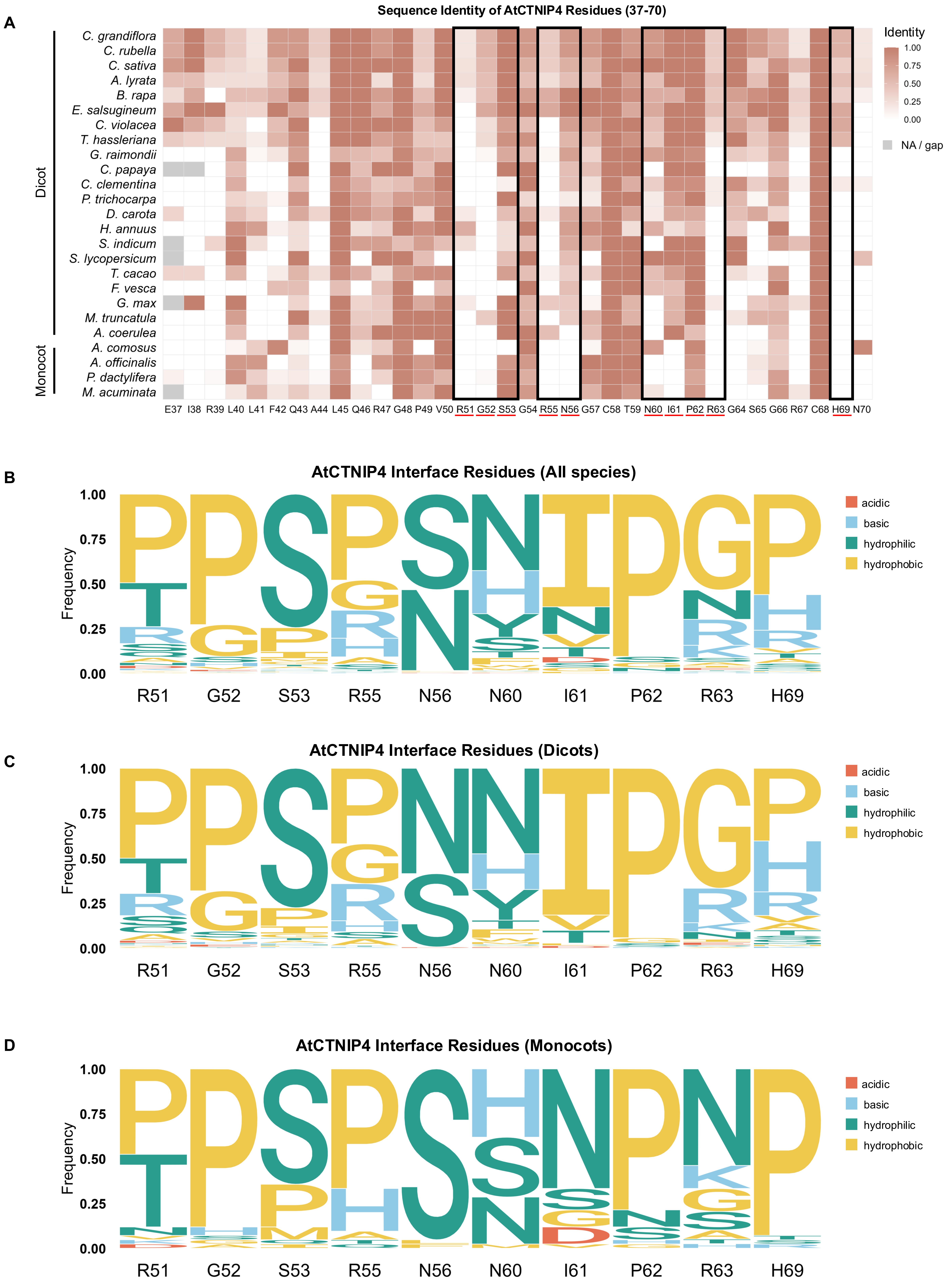

### Supplementary Figure 11

**A**

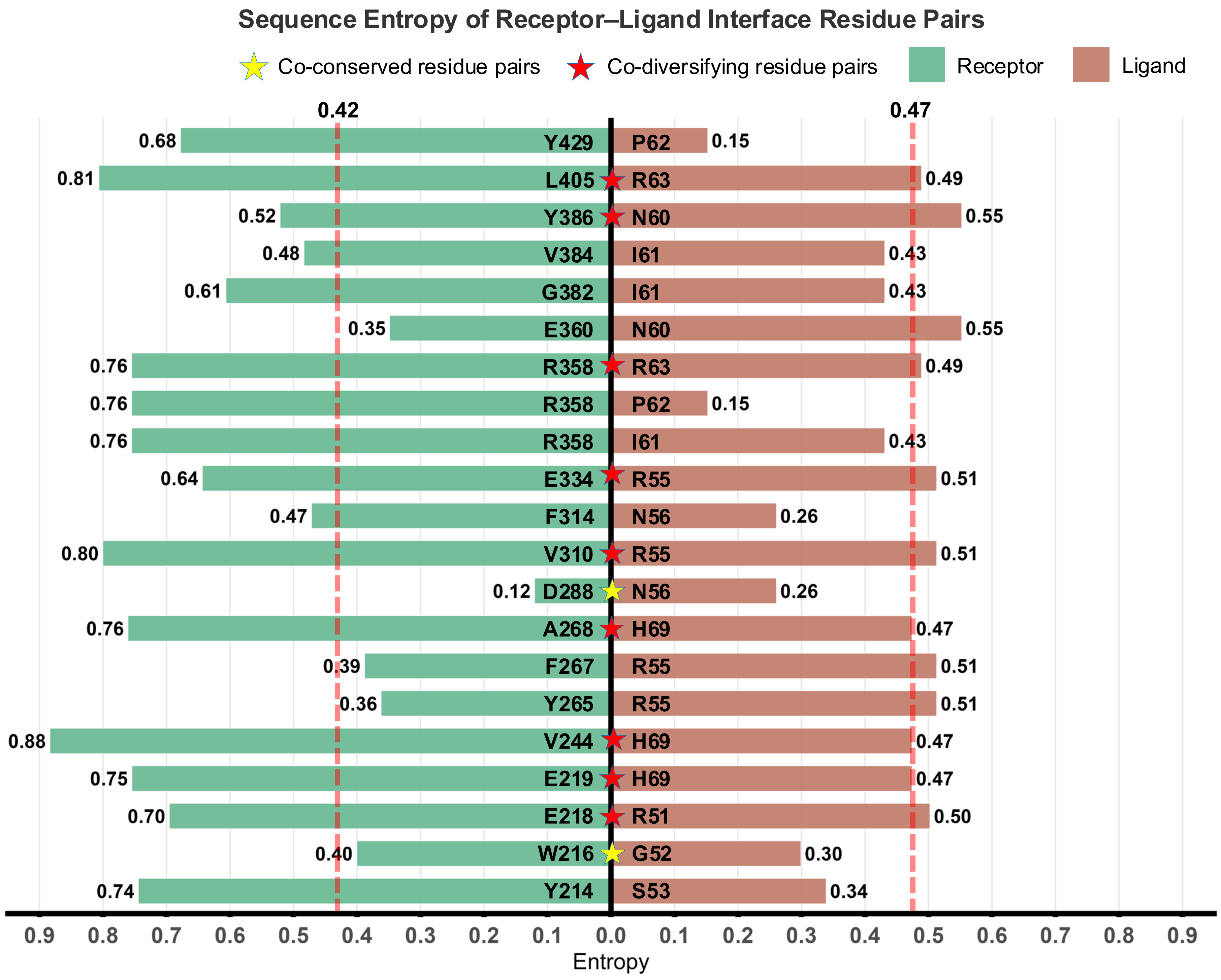

### Supplementary Figure 12

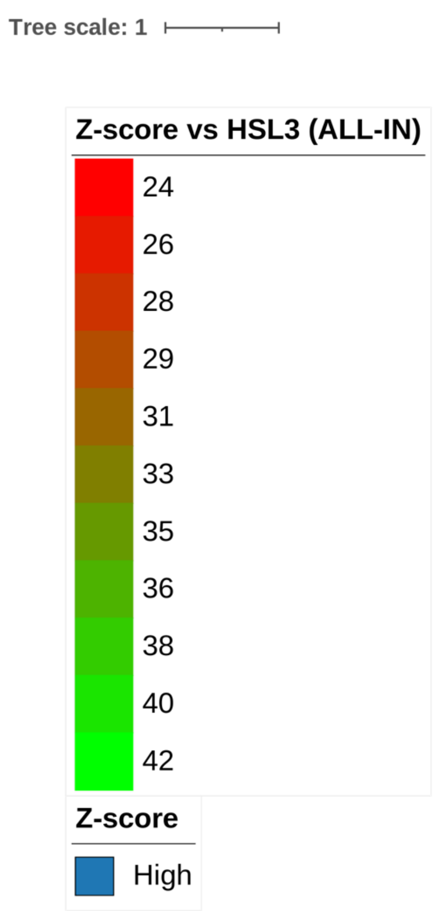


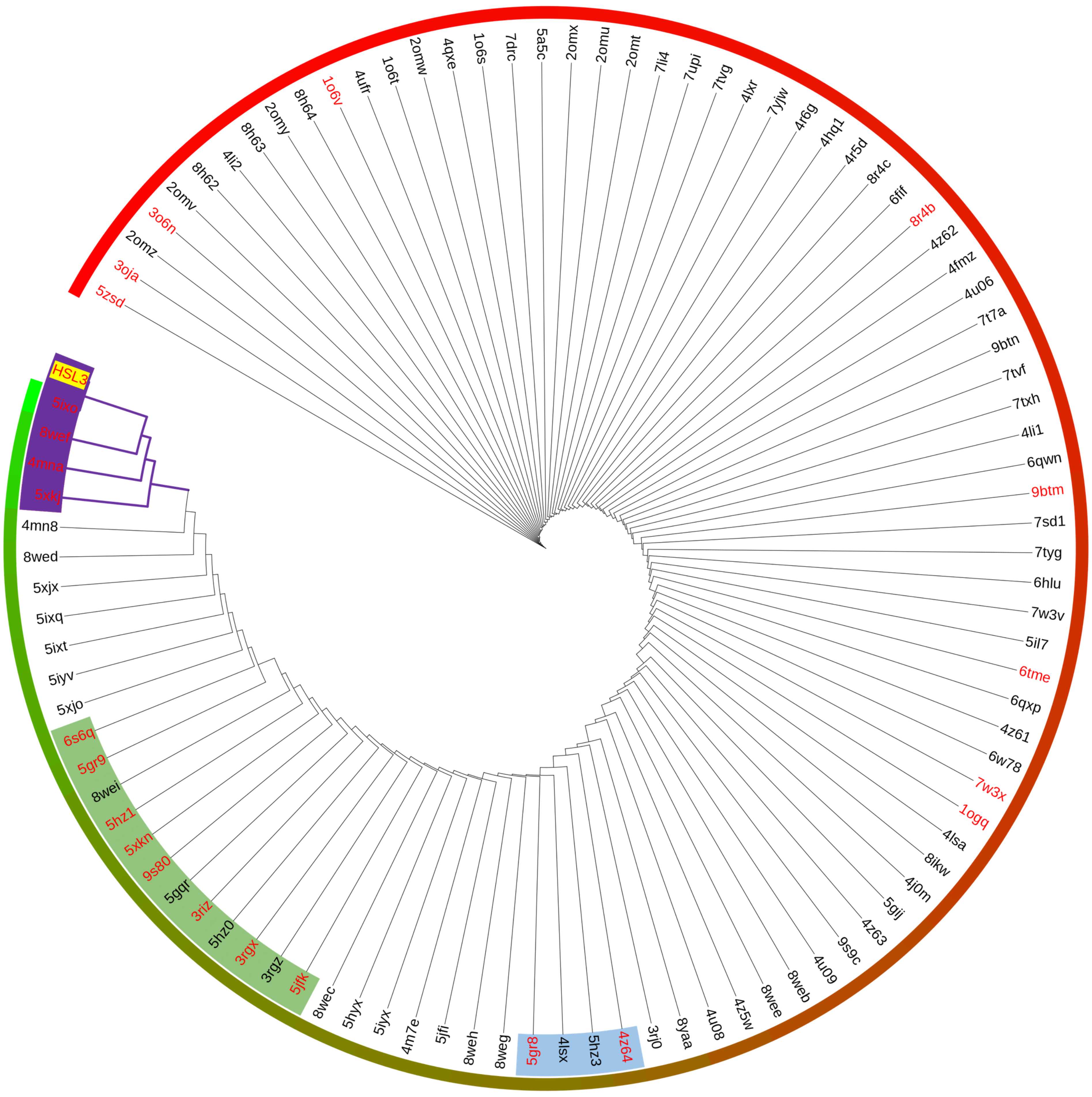


| PDB ID | Protein names |
|---|---|
| 5IXO | *Arabidopsis thaliana* HAESA |
| 8WEF | *Brassica napus* MIK2 |
| 4MNA | *Arabidopsis thaliana* FLS2 |
| 5XKJ | *Arabidopsis thaliana* ERL1 |
| 6S6Q | *Arabidopsis thaliana* GASSHO1/SCHENGEN3 |
| 5GR9 | *Arabidopsis thaliana* PXY-TDIF/CLE41 |
| 5HZ1 | *Arabidopsis thaliana* RGFR1 |
| 5XKN | *Arabidopsis thaliana* ERL2 with EPFL4 |
| 9S80 | *Arabidopsis thaliana* BRL3 with brassinolide |
| 3RIZ | *Arabidopsis thaliana* steroid receptor BRI1 |
| 3RGX | *Arabidopsis thaliana* BRL1 with brassinolide |
| 5JFK | *Arabidopsis thaliana* TDR receptor |
| 5GR8 | *Arabidopsis thaliana* PEPR1-AtPEP1 |
| 4Z64 | *Arabidopsis thaliana* hormone receptor |
| 1OGQ | *Phaseolus vulgaris* PGIP |
| 7W3X | *Nicotiana benthamiana* RXEG1 |
| 6TME | *Arabidopsis thaliana* LRX8 with RALF4 |
| 9BTM | *Homo sapiens* Shoc2 with Nras |
| 8R4B | *Chlorobaculum tepidum* Roco protein |
| 1O6V | *Listeria monocytogenes* EGD-e Internalin |
| 3O6N | *Anopheles gambiae* APL1 |
| 3OJA | *Anopheles gambiae* LRIM/APL1C complex |
| 5ZSD | *Macaca mulatta* TLR7 with IMDQ, GGUUGG |

### Supplementary Table 1

Cryo-EM data collection, refinement, and validation statistics.

| | HSL3 free state | HSL3/CTNIP4 complex |
|---|---|---|
| **PDB entry** | 9XNU | 9XNT |
| **EMDB entry** | 67060 | 67059 |
| **Data collection and processing** | | |
| Magnification | 130,000 | 130,000 |
| Microscope | TFS Krios G4 | TFS Krios G4 |
| Voltage (kV) | 300 | 300 |
| Detector | Gatan K3 BioQuantum | Gatan K3 BioQuantum |
| Energy filter | 20ev slit | 20ev slit |
| Electric exposure ($e^-$/Å) | 65.6 | 61.3 |
| Defocus range (μm) | From -0.7 to -1.8 | From -0.7 to -1.8 |
| Pixel size (Å) | 0.648 | 0.648 |
| Data Processing Program | CryoSPARC (v.4.5.3) | CryoSPARC (v.4.5.3) |
| Movies | 21,554 | 8,404 |
| Initial / Final particle images (no.) | 17,976,959 / 1,740,510 | 7,021,765 / 350,460 |
| Symmetry imposed | C1 | C1 |
| FSC threshold | 0.143 | 0.143 |
| **Refinement** | | |
| Refinement Program | PHENIX (v.1.21.1), Coot (0.9.8.96 EL) | PHENIX (v.1.21.1), Coot (0.9.8.96 EL) |
| Model resolution (Å) | 2.63 | 2.59 |
| FSC threshold | 0.143 | 0.143 |
| Model composition | | |
| Non-hydrogen atoms | 4327 | 4493 |
| Protein residues | 531 | 554 |
| Ligands (NAG) | 13 | 13 |
| Ligands (BMA) | - | 1 |
| RMS deviations | | |
| Bond length (Å) | 0.003 | 0.002 |
| Bond angles (°) | 0.548 | 0.648 |
| Validation | | |
| MolProbity score | 2.53 | 2.82 |
| Clashscore | 13.79 | 16.77 |
| Ramachandran plot | | |
| Favored /Allowed (%) | 90.74 / 9.26 | 90.73 / 9.27 |
| Disallowed (%) | 0 | 0 |
| Mask CC | 0.74 | 0.66 |

### Supplementary Table 2

Comprehensive interaction map of the HSL3–CTNIP4 Binding Interface

| HSL3 residues | Distance (Å) | CTNIP residues |
|---|---|---|
| | Polar contact [a] | |
| Glu 218 [OE2] | 3.52 | Arg 51 [NH2] |
| Glu 219 [OE2] | 3.65 | His 69 [NE2] |
| Asp 288 [OD2] | 3.53 | Asn 56 [N] |
| Glu 334 [OE2] | 3.79 | Arg 55 [NH2] |
| Arg 358 [NH2] | 3.68 | Pro 62 [O] |
| Arg 358 [NH1] | 3.67 | Pro 62 [O] |
| Arg 358 [NH1] | 3.56 | Arg 63 [O] |
| Tyr 386 [OH] | 3.88 | Asn 60 [ND2] |
| | Hydrogen bond | |
| Tyr 214 [OH] | 2.86 | Ser 53 [OG] |
| Arg 358 [NH2] | 3.31 | Ile 61 [O] |
| Glu 360 [OE2] | 3.5 | Asn 60 [ND2] |
| | Salt bridge [b] | |
| Glu 218 [OE2] | 3.52 | Arg 51 [NH2] |
| Glu 219 [OE2] | 3.65 | His 69 [NE2] |
| Glu 334 [OE2] | 3.79 | Arg 55 [NH2] |
| | Non-bonded Van der Waals contacts [c] | |
| Trp 216 [CZ2] | 3.24 | Gly 52 [C] |
| Val 244 [CG2] | 3.56 | His 69 [CD2] |
| Tyr 265 [CE1] | 3.99 | Arg 55 [CG] |
| Phe 267 [CZ] | 3.54 | Arg 55 [C] |
| Ala 268 [CB] | 4.12 | His 69 [CB] |
| Val 310 [CG1] | 3.71 | Arg 55 [CD] |
| Phe 314 [CZ] | 3.97 | Asn 56 [CB] |
| Gly 382 [CA] | 3.81 | Ile 61 [CD1] |
| Val 384 [CG2] | 3.66 | Ile 61 [CG1] |
| Leu 405 [CD1] | 3.71 | Arg 63 [CD] |
| Tyr 429 [CD2] | 3.71 | Pro 62 [CG] |

[a] Polar contacts (3.5–4.0 Å) represent weak donor–acceptor interactions that may become water-mediated hydrogen bonds depending on hydration in higher-resolution maps.
[b] Salt bridges follow PISA classification (electrostatic donor–acceptor ≤ 4.0 Å).
[c] Van der Waals contacts were catalogued using distance thresholds ≤ 4.5 Å.
